# Cell Competition Eliminates Aneuploid Human Pluripotent Stem Cells

**DOI:** 10.1101/2024.05.08.593217

**Authors:** Amanda Ya, Chenhui Deng, Kristina M. Godek

## Abstract

Human pluripotent stem cells (hPSCs) maintain diploid populations for generations despite a persistently high rate of mitotic errors that cause aneuploidy, or chromosome imbalances. Consequently, to maintain genome stability, aneuploidy must inhibit hPSC proliferation, but the mechanisms are unknown. Here, we surprisingly find that homogeneous aneuploid populations of hPSCs proliferate unlike aneuploid non-transformed somatic cells. Instead, in mosaic populations, cell non-autonomous competition between neighboring diploid and aneuploid hPSCs eliminates less fit aneuploid cells. Aneuploid hPSCs with lower Myc or higher p53 levels relative to diploid neighbors are outcompeted but conversely gain a selective advantage when Myc and p53 relative abundance switches. Thus, although hPSCs frequently missegregate chromosomes and inherently tolerate aneuploidy, Myc- and p53-driven cell competition preserves their genome integrity. These findings have important implications for the use of hPSCs in regenerative medicine and for how diploid human embryos are established despite the prevalence of aneuploidy during early development.

## Introduction

Paramount to using human pluripotent stem cells (hPSCs), including embryonic and induced PSCs, in regenerative medicine is the preservation of pluripotency and genomic stability. However, hPSCs often become aneuploid with abnormal segmental and/or whole chromosome copy numbers during prolonged *in vitro* propagation due to culture adaptation that selects for clonal and stable aneuploid karyotypes with a fitness advantage over diploid hPSCs^1–4^. Likewise, aneuploidy also occurs in human preimplantation embryonic cells, with >70% of *in vitro* fertilized (IVF) preimplantation embryos being aneuploid^5–8^ and aneuploidy being the leading cause of miscarriages^9,10^ and congenital birth defects in humans^9,11^.

Mitotic chromosome segregation errors cause segmental and whole chromosome aneuploidies^12,13^, and surprisingly, hPSCs have a persistently high rate of mitotic errors compared to human diploid somatic cells, with lagging chromosomes in anaphase being the most frequent error^14^. In somatic cells, lagging chromosome rates are proportional to chromosome missegregation rates which generates aneuploid progeny^12^ and thus the high rate of lagging chromosomes in hPSCs is likely the leading cause of aneuploidy in these cells. Yet, despite such frequent mitotic errors, hPSCs maintain clonal diploid populations for generations that typically do not become aneuploid until extended passaging^1–4^.

Similar to hPSCs, human preimplantation embryonic cells have a high rate of post-fertilization mitotic errors, including lagging chromosomes, resulting in mitotic errors being the most common cause of aneuploidy in preimplantation embryos^5–8,15–17^. A consequence of mitotic chromosome missegregation is the generation of mosaic embryos composed of aneuploid and diploid cells within a single embryo. Yet, transferred IVF mosaic embryos can produce healthy offspring with diploid karyotypes^18^. Thus, aneuploidy must impose a fitness barrier to hPSC and embryonic cell propagation, but the mechanisms and how these barriers can be overcome are unknown.

Aneuploid somatic cell proliferation is prevented by cell intrinsic consequences including induction of DNA damage^19–22^, DNA replication stress^20–22^, genomic instability^20–22^, p53- and p21-dependent cell cycle arrest^23–25^, and senescence^20^. Yet, prior work suggests that hPSCs are more tolerant toward aneuploidy and do not exhibit a robust intrinsic response until exit from pluripotency upon differentiation^18^. Consequently, this raises the possibility that aneuploid hPSCs in an undifferentiated state are eliminated by alternative pathways. Conversely, previous findings demonstrate that culture adapted aneuploid hPSCs can outcompete diploid hPSCs through mechanical competition mediated by the cytosolic vs. nuclear localization of the transcriptional coactivator yes-associated protein 1 (YAP)^3^. However, the initial molecular events that allow aneuploid hPSCs to propagate and acquire a fitness advantage over diploid hPSCs remain unknown. Here, we investigate both the immediate consequences of inducing aneuploidy in hPSCs and the long-term fate of aneuploid and diploid hPSCs in mosaic populations to address these unanswered questions and shed light on how diploid human embryos might be established during development despite the prevalence of mitotic errors and aneuploidy in preimplantation embryonic cells.

## Results

### hESCs Tolerate Aneuploidy

To study the consequences of aneuploidy in hPSCs, we used the MPS1 kinase inhibitor reversine^26^ to increase mitotic error rates and chromosome missegregation. MPS1 plays roles in both spindle assembly checkpoint signaling to prevent premature anaphase onset and the correction of improper chromosome microtubule attachments to ensure accurate chromosome segregation during mitosis^26^. We constructed H1 human embryonic stem cells (hESCs) that stably express GFP tagged histone H2B (H1 H2B-GFP) to quantify mitotic error rates using time-lapse live-cell imaging. We validated that the protein abundance of the core pluripotency transcription factors Oct4, Sox2, Nanog, and Myc was similar between H1 H2B-GFP and parental H1 hESC populations and that H1 H2B-GFP hESCs do not exhibit any clonal chromosome abnormalities (Figures S1A-C). Crucially, we titrated the dose of reversine to the lowest concentration that significantly increased mitotic errors causing numerical, or whole chromosome missegregation^12^, but not errors causing structural, or segmental abnormalities^13^ (Figures 1A and S1D). Furthermore, there was a marginally significant increase or decrease in prometaphase and metaphase durations, respectively, but overall mitotic timing was unchanged (Figure S1E). Thus, MPS1 activity is partially compromised at this dose since further inhibition with higher concentrations of reversine accelerate mitosis^26^.

**Figure 1.**
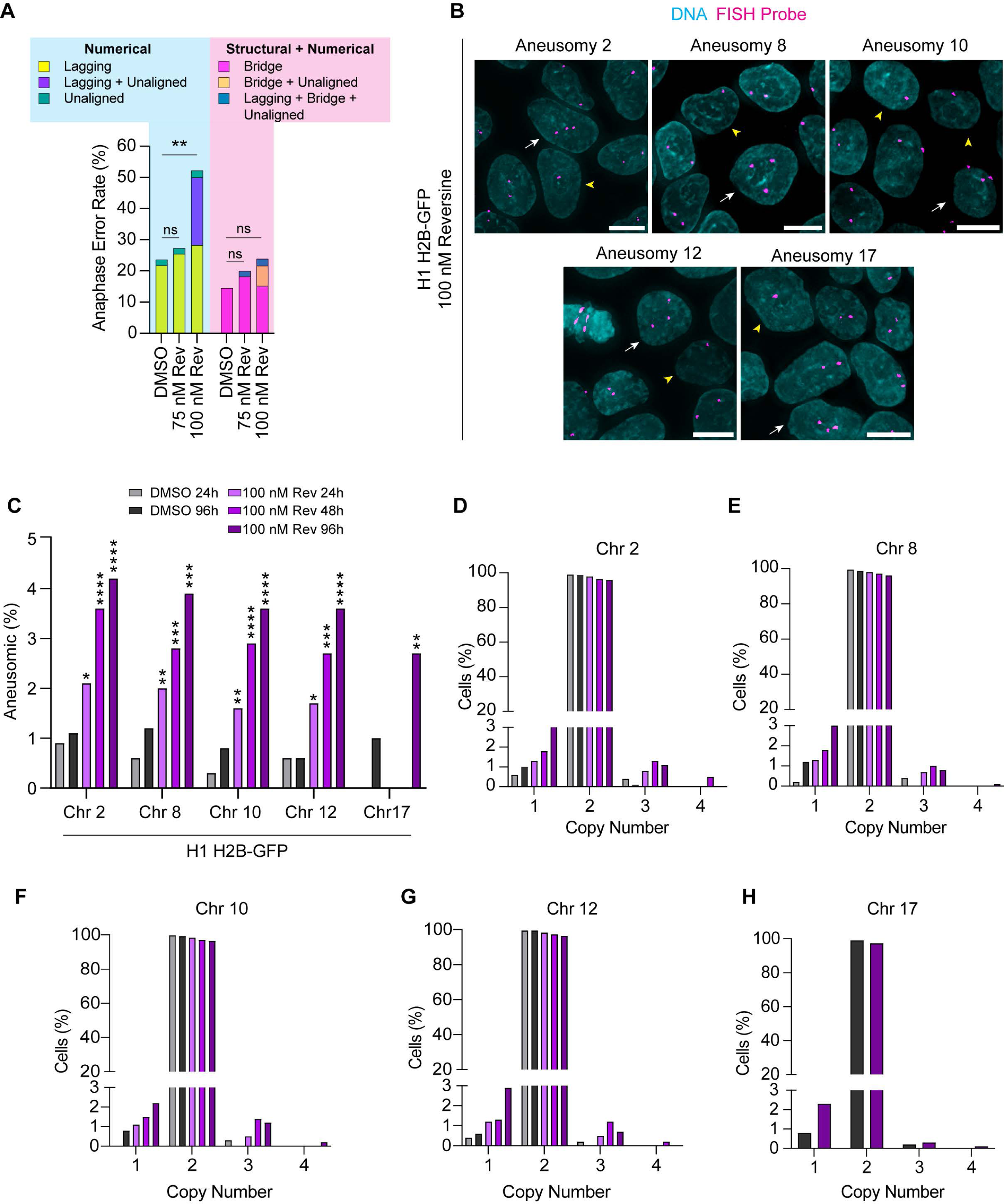
Inhibiting MPS1 kinase increases mitotic errors and aneuploidy in hESCs. (**A**) The percent of anaphase errors quantified from time-lapse live-cell microscopy of H1 H2B- GFP hESCs treated with DMSO, 75 nM, or 100 nM reversine (Rev). Lagging and unaligned chromosomes cause numerical, whole chromosome aneuploidies while chromosome bridges cause structural aneuploidies. n > 45 cells from three independent experiments. (**B**-**H**) The copy numbers of chromosomes 2, 8, 10, 12, or 17 in H1 H2B-GFP hESCs treated with DMSO or 100 nM reversine for 24h, 48h, or 96h determined using FISH. Interphase cells were scored for copy numbers. (**B**) Representative images show aneuploid trisomic (white arrow) or monosomic (yellow arrowhead) cells. Scale bars: 10 μm. (**C**) Aneusomic percent, including gains and losses, of chromosomes 2, 8, 10, 12, and 17. For each chromosome, the aneusomic percent for reversine treated cells was compared to their respective DMSO controls, with 48h reversine compared to 24h DMSO. Copy number distribution of chromosomes 2 (**D**), 8 (**E**), 10 (**F**), 12 (**G**), or 17 (**H**). n = 1000 cells pooled per FISH probe from two independent experiments. ns p > 0.05, *p < 0.05, **p < 0.01, ***p < 0.001, and ****p < 0.0001 using a two-tailed Fisher’s exact test (**A** and **C**). See also Figure S1.

To assess whether partial MPS1 inhibition caused a corresponding increase in chromosome missegregation and aneuploidy, we determined individual chromosome copy numbers in single H1 H2B-GFP hESCs using fluorescence *in situ* hybridization (FISH) and quantified the percent of aneusomic cells that deviated from the diploid two copies. The percent of aneuploid H1 H2B-GFP hESCs significantly increased with the duration of reversine treatment due to both chromosome gains and losses compared to DMSO controls (Figures 1B-H). Of note, fresh media with 100 nM reversine or DMSO was added daily during the treatment time course. Furthermore, we observed no significant difference in the aneusomic rate (Fisher’s exact test p > 0.05) among chromosomes 2, 8, 10, 12, or 17 unlike in somatic cells^27^ (Figure 1C) demonstrating that chromosome missegregation was random in H1 H2B-GFP hESCs.

Moreover, given that the aneusomic frequency was ∼3-4% per chromosome after 96 hr reversine treatment, we estimate that nearly all cells in the population become aneuploid when accounting for all 46 chromosomes hence generating a homogeneous population of aneuploid hESCs with diverse karyotypes.

Our observation that H1 H2B-GFP hESCs seemed to tolerate chromosome missegregation and aneuploidy was surprising given that in somatic cells chromosome missegregation rapidly induces cell autonomous responses that prevent the proliferation of aneuploid progenies^20,21,23–25^. To test whether induction of aneuploidy inhibited hESC proliferation, we measured the growth of H1 H2B-GFP hESCs in 100 nM reversine over the course of 96 hrs. We included somatic diploid RPE-1 epithelial cells expressing H2B-GFP as a positive control (Figure 2A). As expected, 100 nM reversine treatment elevated the numerical and structural error rates without appreciably changing mitotic duration in RPE-1 H2B-GFP cells (Figure S2A). This significantly increased chromosome missegregation and aneuploidy (Figures S2B and S2C). In contrast to hESCs, but in agreement with previous findings^27^, there was biased chromosome missegregation in RPE-1 H2B-GFP cells with significantly increased missegregation of chromosome 2 compared to the other chromosomes (aneusomic rate chromosome 2 vs. other chromosomes Fisher’s exact test p < 0.05) (Figure S2C). Moreover, the induction of aneuploidy in RPE-1 H2B-GFP cells significantly increased p53 and p21 levels (Figures 2C and 2D) and inhibited their proliferation (Figure 2B) compared to DMSO controls. Thus, as expected^23,24^, aneuploidy in somatic cells causes cell autonomous responses that prevent the propagation of aneuploid cells.

**Figure 2.**
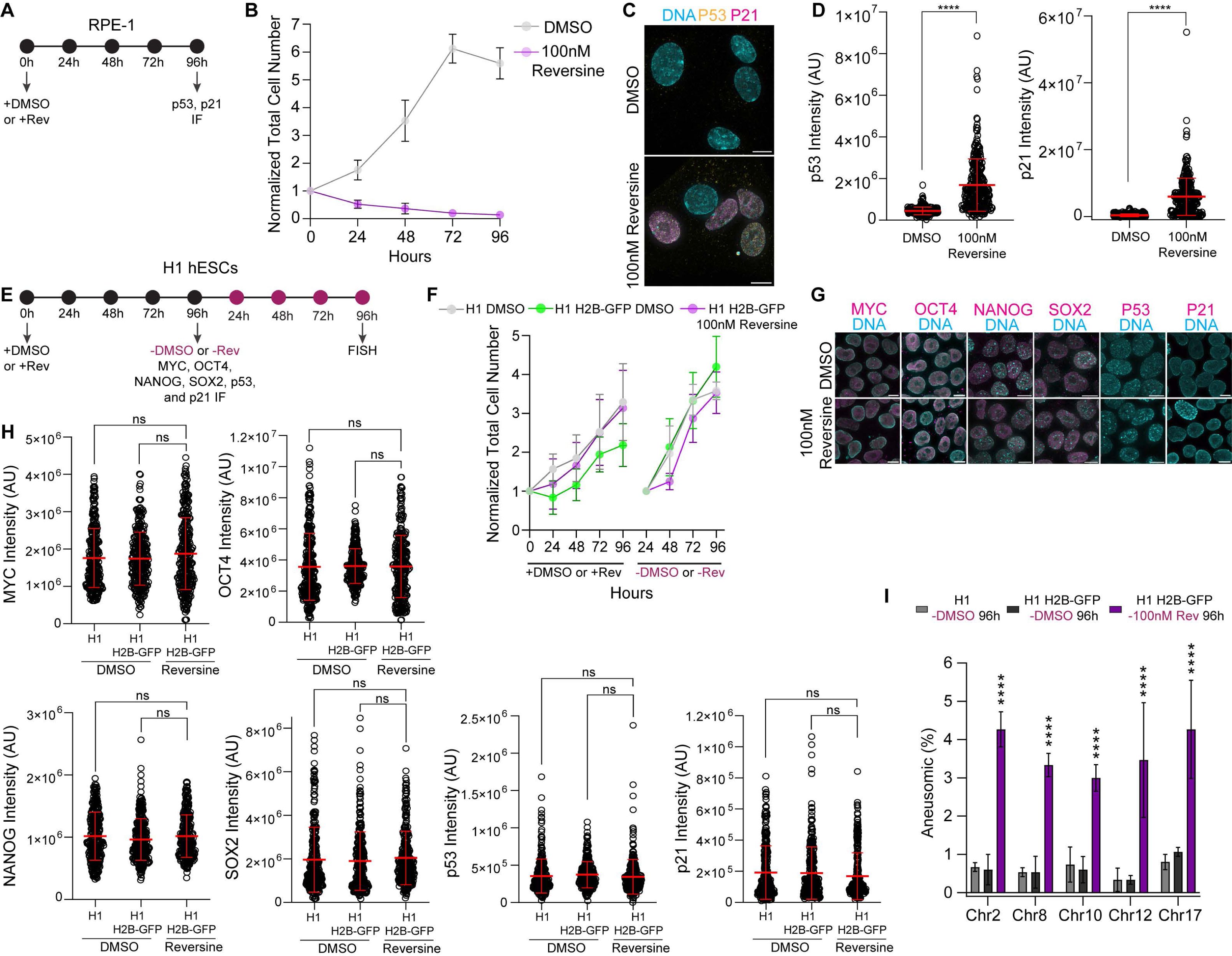
hESCs tolerate aneuploidy. (**A**) Experimental design testing the aneuploidy tolerance of somatic RPE-1 cells. (**B**-**D**) RPE-1 cells were treated with DMSO or 100 nM reversine for 96h to induce aneuploidy. (**B**) Growth curves of diploid vs. aneuploid RPE-1 cells. Each time point is normalized to 0h. Mean ± SD (**C**) Representative IF images of diploid vs. aneuploid RPE-1 cells. Shown is DNA (cyan), p53 (yellow), and p21 (magenta). Scale bars: 10 μm. (**D**) Quantification of p53 and p21 protein levels by IF in diploid vs. aneuploid RPE-1 cells. n = 300 cells; mean ± SD. (**E**) Experimental design testing the aneuploidy tolerance of H1 hESCs. (**F**-**I**) H1 and H1 H2B-GFP hESCs were treated with DMSO or 100 nM reversine for 96h to induce aneuploidy. (**F**) Growth of curves of diploid H1 and H1 H2B-GFP hESCs vs. aneuploid H1 H2B-GFP hESCs. Each time point is normalized to 0h. Mean ± SD (**G**) Representative IF images of diploid and aneuploid H1 H2B-GFP hESCs. Shown is DNA (cyan) and Myc, Oct4, Nanog, Sox2, p53, or p21 (magenta). Scale bars: 10 μm. (**H**) Quantification of Myc, Oct4, Nanog, Sox2, p53, or p21 protein levels by IF in diploid H1 and H1 H2B-GFP hESCs vs. aneuploid H1 H2B-GFP hESCs. n = 300 cells; mean ± SD. (**I**) Aneusomic percent, including gains and losses, determined using FISH for chromosomes 2, 8, 10, 12, or 17 in H1 and H2B-GFP hESCs upon withdrawal of DMSO or reversine for 96h. For each chromosome, the aneusomic percent of their respective DMSO and reversine treated cells were compared. n = 1500 cells per FISH probe; mean± SD. Three independent experiments (**B**, **D**, **F**, **H**, and **I**); ns p > 0.05 and ****p < 0.0001 using a two-tailed Student’s t-test (**D**); a one-way ANOVA with Tukey’s correction (**H**), or a two-tailed Fisher’s exact test (**I**). See also Figure S2.

In contrast, H1 H2B-GFP hESCs proliferated upon continuous induction of aneuploidy for 96 hrs comparable to the growth of DMSO controls (Figures 2E and 2F). Moreover, aneuploidy did not induce a significant increase of p53 or p21 in H1 H2B-GFP hESCs, unlike in RPE-1 somatic cells (Figures 2D, 2G, and 2H). One possibility is that aneuploid H1 H2B-GFP hESCs proliferated due to the clonal acquisition of mutations in p53. In somatic and cancer cells, p53 deletion or mutation permits the survival of aneuploid cells^24,28^, and hESCs frequently acquire p53 mutations during *in vitro* propagation^29,30^. We tested for wild-type p53 functionality by treating H1 H2B-GFP hESCs with the DNA damaging agent doxorubicin. DNA damage induced similar p53 levels but attenuated p21 levels in H1 H2B-GFP hESCs compared to RPE- 1 somatic cells in agreement with previous reports^31–34^ (Figures S2D-G). Also, we measured the survival of H1 H2B-GFP hESCs after 24 hr doxorubicin treatment because p53 mutations confer resistance to apoptosis upon DNA damage induced by doxorubicin^35^. H1 H2B-GFP hESCs all died by 24 hrs in the presence of doxorubicin (Figure S2H). Thus, p53 exhibits wild-type function in H1 H2B-GFP hESCs and a clonal p53 mutation is unlikely to account for the survival of aneuploid hESCs.

We also considered that inhibiting MPS1 kinase using reversine not only increases chromosome missegregation but also permits the survival of aneuploid hESCs. To test this, we passaged aneuploid H1 H2B-GFP hESCs following reversine withdrawal for 96 hrs and measured their proliferation and the frequency of aneusomic cells (Figure 2E). H1 H2B-GFP hESCs proliferated after reversine withdrawal comparable to DMSO controls (Figure 2F). Notably, the aneusomic frequency also remained elevated demonstrating the continued survival and proliferation of aneuploid H1 H2B-GFP hESCs in the absence of reversine (Figure 2I). Lastly, we monitored aneuploid H1 H2B-GFP hESCs for signs of premature differentiation, but there was no significant difference in the abundance of the pluripotency transcription factors Myc, Oct4, Nanog, or Sox2 compared to DMSO control cells (Figure 2H). Thus, hESCs maintain an undifferentiated state upon acute induction of aneuploidy in agreement with previous findings^18^. Collectively, these results demonstrate that in contrast to somatic cells, homogeneous aneuploid populations of hESCs are inherently tolerant of aneuploidy.

### Cell Competition Occurs between Aneuploid and Diploid hESCs in Mosaic Populations

Given the lack of a rapid and robust intrinsic response to aneuploidy in hESCs, we explored cell non-autonomous competition as a mechanism that eliminates aneuploid hESCs, particularly since aneuploid cells are at least initially in a mosaic environment with diploid cells during routine culturing. Cell competition is characterized by (1) distinct cell populations exhibiting different growth rates in a mosaic environment, (2) the elimination of less fit, but viable, “loser” cells due to local interactions with neighboring more fit “winner” cells, and (3) the potential for “loser” or “winner” status and cell fate to switch because these phenotypes are dependent on the relative fitness of neighboring cells^36^. To investigate competition between aneuploid and diploid hESCs, we measured the proportion of aneuploid and diploid cells in unperturbed mosaic populations over time using two complementary methods: (1) G-banding karyotyping to monitor the copy number of all chromosomes simultaneously in single cells and (2) FISH using chromosome specific probes. We compared mosaic CSES8 and CSES22 hESCs with aneuploid karyotypes of different origins and control clonal diploid CSES7 hESCs (Figure 3A). Importantly, all three hESC lines were cultured in identical undifferentiated conditions.

**Figure 3.**
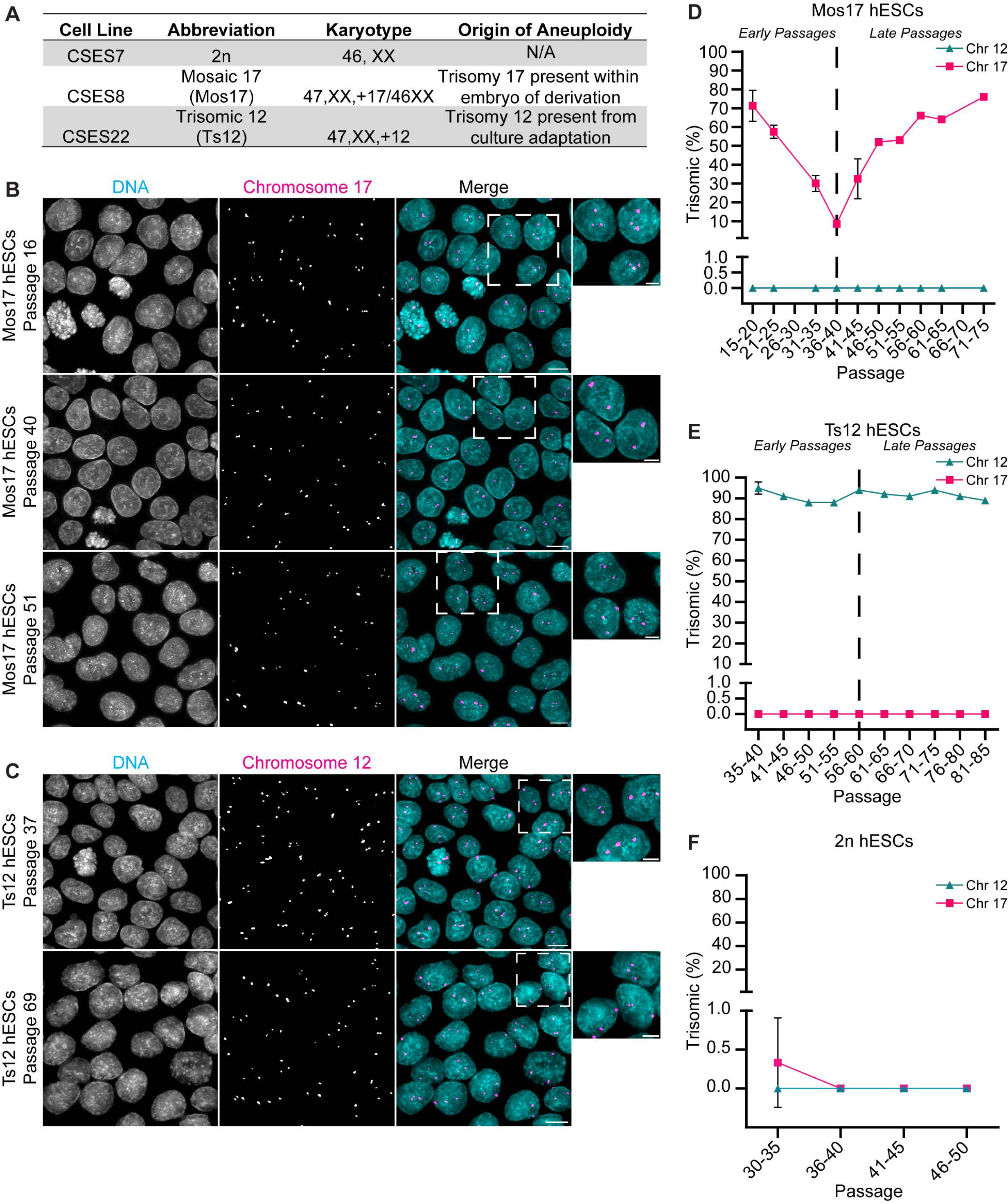
Cell competition between diploid and aneuploid hESCs in mosaic populations. (**A**) Karyotype and origin of aneuploidy in CSES7 (2n), CSES8 (Mos17), and CSES22 (Ts12) hESCs. (**B**-**C**) Representative images of chromosome 17 copy numbers in Mos17 hESCs (**B**) or of chromosome 12 copy numbers in Ts12 hESCs (**C**) from FISH performed at different passages. Shown is DNA (cyan) and chromosome 17 or 12 (magenta). Insets highlight disomic or trisomic chromosome 17 or 12 cells. Scale bars: 10 μm or 5 μm for insets. (**D**-**F**) Percent of trisomic chromosome 17 (magenta) or 12 (cyan) Mos17 (**D**), Ts12 (**E**), or 2n (**F**) hESCs during growth in culture. Dashed line separates passages categorized as “early” or “late.” Data points are binned into groups with 5 passages. For each data point a minimum of two experiments performed. n = 100 cells per replicate; mean ± SD. Note for some data points the error bars are smaller than the data symbol. See also Figure S3.

CSES7 hESCs retained a clonal diploid population during extended propagation as shown previously^37^ (Figures 3F, S3B, and S3F-G). Twenty of twenty cells analyzed by G-banding karyotyping at passages 36 and 53 were diploid (Figure S3B). Thus, CSES7 hESCs (here after referred to as 2n) exhibited no appreciable aneuploid population or mosaicism.

Prior work showed that CSES22 hESCs acquired a clonal abnormality of chromosome 12 trisomy during passaging which outcompeted diploid cells^38^. We confirmed that ∼90% of CSES22 hESCs were trisomic and ∼10% disomic for chromosome 12 during prolonged passaging using FISH (Figures 3C and 3E). Moreover, G-banding karyotyping of passage 53 cells identified a single cell disomic for chromosome 12 (Figure S3C) supporting our FISH results (Figure 3E). HPSCs persistently commit mitotic errors^14^, so one possible explanation is that random chromosome missegregation sporadically causes loss of a copy of chromosome 12 producing disomic progenies. More importantly, however, the relative proportion of trisomic vs. disomic chromosome 12 cells was consistent over time indicating that trisomic chromosome 12 cells retained a competitive advantage over disomic cells (Figure 3E). Thus, CSES22 hESCs (here after referred to as Ts12) exhibited a low level of mosaicism but were a predominately stable and clonal aneuploid population that arose due to culture adaptation and the gain of a selective advantage over diploid cells.

CSES8 hESCs were derived from an IVF embryo with chromosome 17 trisomy^39^. Although CSES8 hESCs were previously characterized as uniformly trisomic for chromosome 17^2,39^, we observed that CSES8 hESCs (here after referred to as Mos17) were a mosaic population of disomic and trisomic chromosome 17 cells and that the proportion of trisomic vs. disomic chromosome 17 cells in the population changed over time (Figures 3B, 3D, S3D-E, and S3H). Importantly, short-tandem repeat profiling demonstrated that other contaminating cells are not the source of mosaicism (Figure S3A). A possible explanation is that the embryo from which the Mos17 hESCs were derived was composed of both disomic and trisomic chromosome 17 embryonic cells; however, it is not feasible to test this.

At the earliest passage available to us, the Mos17 hESCs were ∼75% trisomic and ∼25% disomic for chromosome 17 (Figures 3B and 3D). Initially, the proportion of trisomic chromosome 17 cells declined in the population over a period of weeks reaching a low point of ∼10% (Figures 3D, S3D, and S3H). One possibility is that Mos17 hESCs had an elevated basal rate of genomic instability resulting in frequent but random chromosome missegregation producing chromosome 17 disomic cells. To determine if Mos17 hESCs exhibited elevated genomic instability, we karyotyped Mos17 hESCs at passages 20 and 30 and monitored the copy number of chromosome 12 in the population using FISH but detected no other clonal abnormalities (Figures 3D and S3D). Alternatively, a single copy of chromosome 17 could be iteratively lost in Mos17 hESCs. This is highly improbable, however, because it requires repeated and selective chromosome 17 missegregation, and we found no evidence to support this in hESCs (Figure 1C). Thus, the possibility that genomic instability explains the observed population dynamics of trisomic vs. disomic cells is unlikely. Accordingly, aneuploid hESCs with trisomy 17 were initially at a selective disadvantage relative to diploid hESCs in the population and were outcompeted by an alternative mechanism. We designated this phase as “early passages” and defined it as ≤ 40 passages for Mos17 hESCs. This corresponds to 25 passages from the starting passage available to us. An equivalent time frame was used to define ≤ 60 as “early passages” for Ts12 hESCs (Figure 3E).

Strikingly, upon further propagation, there was a switch in the proportion of trisomic vs. disomic chromosome 17 cells in the population. Trisomic chromosome 17 cells increased and outcompeted disomic chromosome 17 cells (Figures 3D, S3D, and S3H). Furthermore, karyotype analysis at passage 69 and FISH for chromosome 12 copy numbers demonstrated no other clonal abnormalities indicating that genomic instability likely does not explain the increase in trisomic chromosome 17 cells (Figures 3D and S3D). Thus, aneuploid trisomic 17 hESCs gained a fitness advantage relative to diploid hESCs during this period. We designated this phase as “late passages” and defined it as > 40 passages for Mos17 hESCs with the corresponding time frame in Ts12 hESCs being > 60 passages (Figure 3E). Crucially, we confirmed that the fate of trisomic chromosome 17 cells relative to disomic cells during extended passaging is reproducible (Figure S3H). Collectively, these results demonstrate that in a mosaic population, diploid and aneuploid hESCs exhibit different growth rates with aneuploid hESCs initially being less fit, slower proliferating cells relative to diploid hESCs. Consequently, aneuploid hESCs are outcompeted from the population. However, during extended propagation, the cell fitness of aneuploid relative to diploid hESCs can switch from a selective disadvantage to an advantage with aneuploid hESCs overtaking the population. Thus, meeting key criteria that define cell competition.

### Myc and p53 Mediate Cell Competition Between Aneuploid and Diploid hESCs

Prior studies of embryogenesis and embryonic stem cells (mESCs) in mice demonstrate that a ∼2-fold relative increase in abundance of the transcription factor Myc in a cell compared to its adjacent neighbors confers a fitness advantage and “winner” status while neighbors with relatively lower Myc levels are less fit “loser” cells that are eliminated by apoptosis^36,40–42^. If Myc similarly mediates cell competition between aneuploid and diploid hESCs, this predicts that the relative protein abundance of Myc between neighboring aneuploid and diploid cells determines their cell fitness and fate. To test this, we quantified Myc levels between trisomic and disomic Mos17 or Ts12 hESC neighbors during both “early” and “late” passages by combining standard immunofluorescence using an antibody that recognizes c- and n-Myc with FISH (IF-FISH) (Figures 4A-D). Specifically, we calculated the ratio of Myc intensity between an index trisomic cell and its respective disomic or trisomic adjacent neighbors (Figures 4A-D). We did this for both Mos17 and Ts12 hESC populations. A ratio of 1 indicates equivalent Myc levels between the index trisomic cell and its neighbors. Of note, for these and subsequent experiments, index trisomic cells were chosen solely based upon their chromosome copy number to minimize selection bias. Also, replicates were performed at three different “early” or “late” passage time points.

**Figure 4.**
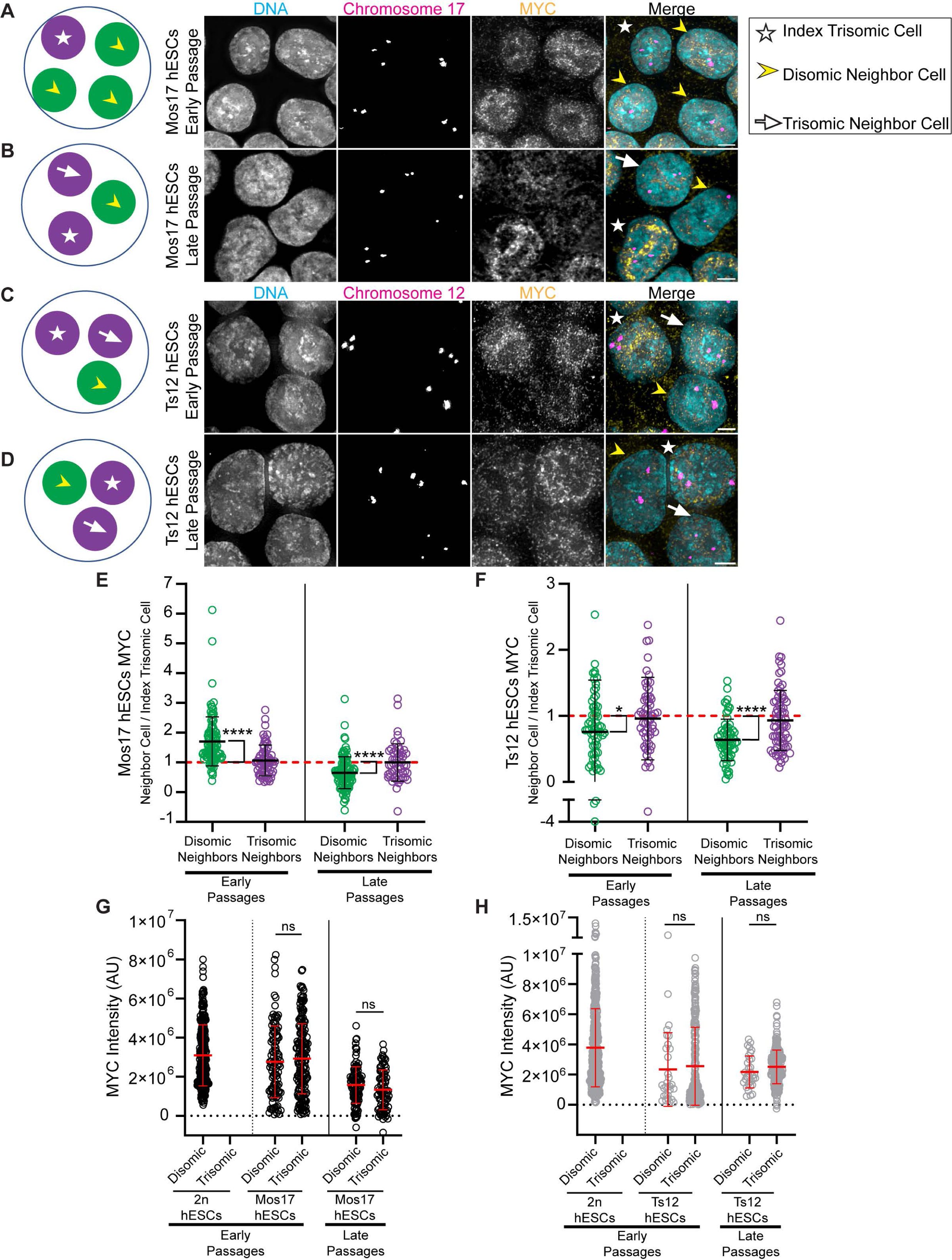
Myc mediates cell competition between aneuploid and diploid hESCs. (**A**-**D**) Cartoon representations (left panels) showing Mos17 (**A** and **B**) or Ts12 (**C** and **D**) hESC neighborhoods comprised of an index trisomic cell (purple, white star), disomic neighbor cells (green, yellow arrowhead), or trisomic neighbor cells (purple, white arrow). Representative IF- FISH images (right panels) of Mos17 hESC neighborhoods at early (**A**) or late (**B**) passage or Ts12 hESC neighborhoods at early (**C**) or late (**D**) passage. Shown is DNA (cyan), chromosome 17 or 12 (magenta), and Myc (yellow). Scale bar: 5 μm. (**E**-**F**) Quantification of Myc protein levels by IF-FISH. Neighbor cells were normalized to index trisomic cells in early (left) and late (right) passage Mos17 (**E**) or Ts12 (**F**) hESCs. Red dashed line indicates a ratio of 1 and equivalent Myc levels between index trisomic cells and neighbors. n = 60 index trisomic cells with > 124 neighbors; mean ± SD. (**G**-**H**) Quantification of Myc protein levels by IF-FISH in randomly selected, non-neighboring 2n and Mos17 hESCs (**G**) or 2n and Ts12 hESCs (**H**) at early (left panels) or late (right panels) passages. n= 300 total cells; mean ± SD. Three independent experiments for both early and late passages (**E**-**H**); ns p > 0.05, *p < 0.05, and ****p < 0.0001 using a one sample t-test compared to hypothetical mean value 1 (**E** and **F**) or a two-tailed Student’s t-test within each cell line for early or late passage (**G** and **H**). See also Figure S4.

Our analysis demonstrated that there was a significant increase in relative Myc levels for disomic Mos17 neighbors compared to index trisomic cells at “early” passages corresponding to when trisomic chromosome 17 cells were less fit and outcompeted. Furthermore, the relative abundance of Myc switched and decreased for disomic neighbors compared to index trisomic Mos17 cells during “late” passages corresponding to when trisomic chromosome 17 cells outcompeted disomic cells (Figure 4E). Moreover, there was no change in relative Myc levels between trisomic neighbors compared to index trisomic cells during either phase demonstrating that abundance differences were exclusive between disomic cells neighboring trisomic cells (Figure 4E). Since gene and protein expression tend to scale with chromosome copy number^43–45^, we also determined that the switch in relative Myc abundance was not due to trisomic or disomic chromosome 17 cells gaining or losing, respectively, copies of chromosome 8 that includes the *Myc* gene locus (Figures S4C and S4E). Finally, for Ts12 hESCs, there was a significant decrease in relative Myc levels for disomic Ts12 neighbors irrespective of “early” or “late” passages (Figure 4F) correlating with the fitness advantage trisomic chromosome 12 cells maintain over disomic cells, and this was not due to alterations in the copy number of chromosome 8 (Figures S4D and S4F). Importantly, there was no significant difference in Myc levels between trisomic vs. disomic Mos17 or Ts12 hESCs at “early” or “late” passages between randomly selected non-neighboring trisomic or disomic cells (Figures 4G, 4H, S4A, and S4B) demonstrating that the absolute levels of Myc do not correlate with cell fitness.

In addition to Myc, studies in mouse embryos and mESCs demonstrate that the relative levels of the transcription factor p53 between neighboring cells determines their competitive ability^42,46–49^. Higher relative levels of wild-type p53 are associated with reduced cell fitness and induction of apoptosis^42,46,48,49^. Thus, we predict that the relative levels of p53 between aneuploid and diploid hESC neighbors will inversely correlate with their cell fitness and fate. To test this prediction, we quantified p53 levels between trisomic and disomic Mos17 or Ts12 hESC neighbors during both “early” and “late” passages using IF-FISH (Figures 5A-D). In support, there was a significant decrease in relative p53 levels for disomic Mos17 neighbors compared to index trisomic cells at “early” passages when trisomic chromosome 17 cells were outcompeted (Figure 5E). However, the *p53* gene locus is located on chromosome 17, and there was a significant increase in absolute p53 levels for trisomic chromosome 17 cells (Figures 5G and S5A). Yet, irrespective of p53 absolute levels, at “late” passages when trisomic chromosome 17 cells gained a fitness advantage, there was no longer a significant difference in the relative abundance of p53 between disomic neighbors and index trisomic Mos17 cells (Figures 5E, 5G and S5A). Thus, the relative abundance of p53 between trisomic and disomic neighbors changed corresponding with a switch in competitive advantage while the absolute levels did not. Moreover, disomic Ts12 hESCs had a significant increase in relative p53 levels at both “early” and “late” passages compared to index trisomic cells (Figure 5F), and the absolute levels of p53 did not correlate with the competitive advantage of trisomic chromosome 12 cells (Figures 5H and S5C).

**Figure 5.**
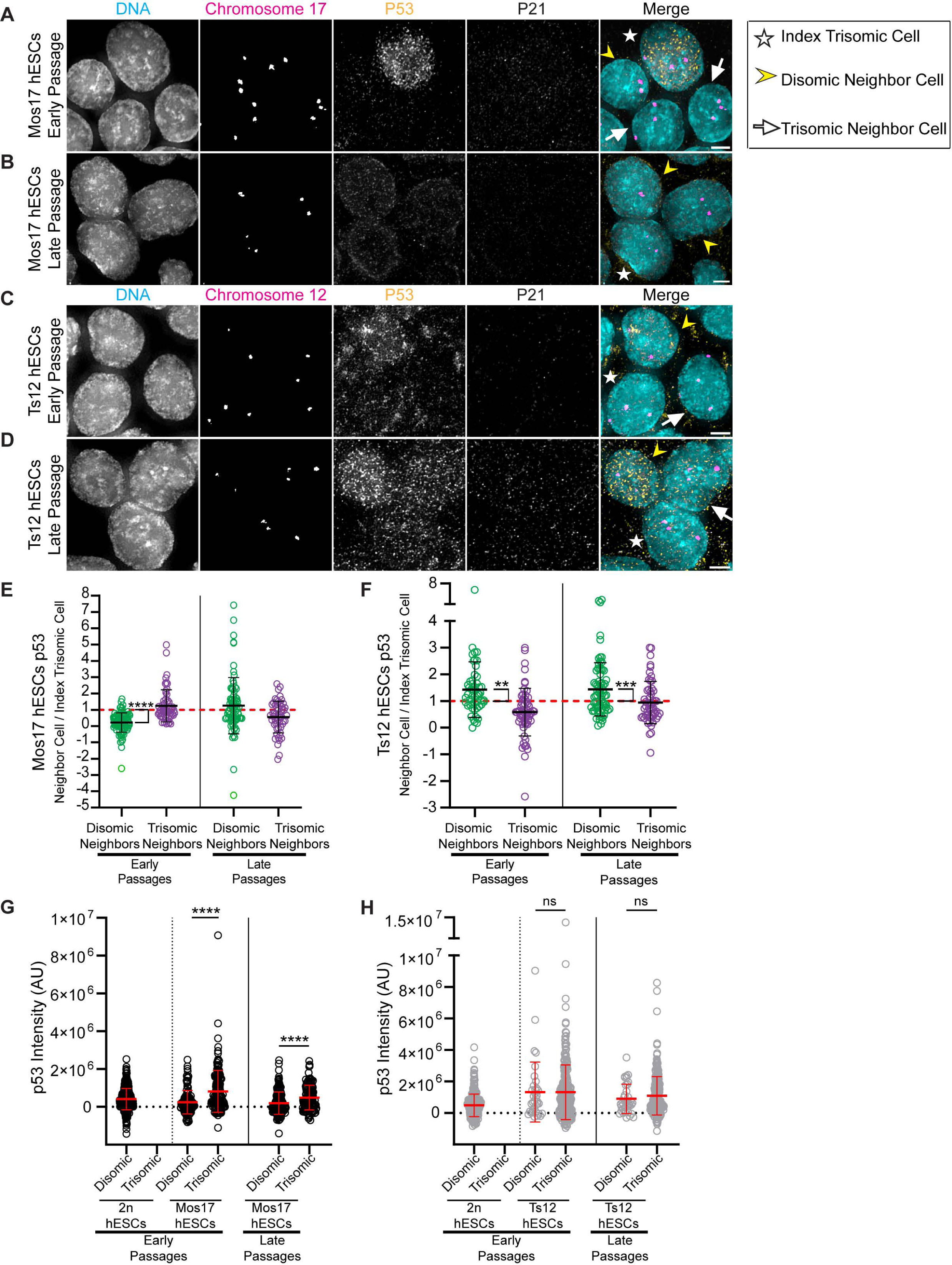
p53 mediates cell competition between aneuploid and diploid hESCs. (**A**-**D**) Representative IF-FISH images of Mos17 early (**A**) or late (**B**) passage or Ts12 early (**C**) or late (**D**) passage hESC neighborhoods comprised of an index trisomic cell (white star), disomic neighbor cells (yellow arrowhead), or trisomic neighbor cells (white arrow). Shown is DNA (cyan), chromosome 17 or 12 (magenta), p53 (yellow), and p21 (gray). Scale bar: 5 μm. (**E**-**F**) Quantification of p53 protein levels by IF-FISH. Neighbor cells were normalized to index trisomic cells in early (left) and late (right) passage Mos17 (**E**) or Ts12 (**F**) hESCs. Red dashed line indicates a ratio of 1 and equivalent p53 levels between index trisomic cells and neighbors. n = 60 index trisomic cells with > 100 neighbors; mean ± SD. (**G**-**H**) Quantification of p53 protein levels by IF-FISH in randomly selected, non-neighboring 2n and Mos17 hESCs (**G**) or 2n and Ts12 hESCs (**H**) at early (left panels) and late (right panels) passages. n= 300 total cells; mean ± SD. Three independent experiments for both early and late passages (**E**-**H**); ns p > 0.05, **p < 0.01, ***p < 0.001, and ****p < 0.0001 using a one sample t-test compared to hypothetical mean value 1 (**E** and **F**) or a two-tailed Student’s t-test within each cell line for early or late passage (**G** and **H**). See also Figure S5.

A well-established downstream target of p53 is cyclin dependent kinase inhibitor p21^50^, and we tested whether the relative levels of p21 correlated with trisomic vs. disomic cell fitness. However, there was no significant difference in relative abundance of p21 between trisomic or disomic Mos17 or Ts12 neighbors at either “early” or “late” passages (Figures 5A-D, S5E, and S5G). Moreover, we observed that both trisomic and disomic Mos17 or Ts12 hESCs expressed low absolute levels of p21 (Figures S5A, S5C, S5F, and S5H), but that DNA damage induced the expected p53 and p21 response in trisomic and disomic Mos17 or Ts12 hESCs^31,33,34^ (Figures S5B, S5D, S5I, and S5J). Thus, p53 and p21 exhibit wild-type functionality, but p53 is acting through mechanisms other than p21 to mediate cell compeition. Combined, these results indicate that relative differences in Myc and p53 abundance between neighboring hESCs drives cell competition between aneuploid and diploid hESCs. Consequently, aneuploid hESCs with relatively lower Myc and higher p53 levels are outcompeted in early passages, but conversely, aneuploid hESCs with relatively higher Myc and lower p53 levels gain a competitive advantage leading to their selection in late passages.

### Competitive Interactions Between Aneuploid and Diploid hESCs Occur in an Undifferentiated State

In mouse embryos and mESCs, Myc and p53 eliminate unfit cells through competitive interactions coinciding with the initiation of differentiation^40,42,46,47^. Although we maintained all hESCs in undifferentiated conditions, a possible consequence of having an aneuploid genome with chromosome imbalances could be loss of an undifferentiated state leading to Myc- and p53-driven cell competition and the elimination of aneuploid hESCs. To test this possibility, we used IF-FISH to quantify the relative levels of the pluripotency transcription factors Oct4, Sox2, and Nanog between trisomic and disomic Mos17 or Ts12 neighbors during both “early” and “late” passages reasoning that decreased expression of these transcription factors indicates premature differentiation. There was no significant difference in the relative levels of Oct4, Sox2, or Nanog between either index trisomic Mos17 or Ts12 hESCs and their respective disomic or trisomic neighbors at “early” or “late” passages (Figures 6A-D, S6E, and S6F). Furthermore, although there were some minor differences in absolute levels of Oct4, Sox2, or Nanog for disomic vs. trisomic Mos17 hESCs, the absolute differences in abundance did not correlate with cell fitness, and there were no significant differences in absolute levels for Ts12 hESCs (Figures 6E, 6F, and S6A-D). These results, in combination with our findings upon acute induction of aneuploidy (Figure 2H), suggest that differentiation of aneuploid hESCs is not responsible for the competitive interactions between aneuploid and diploid hESCs. Rather, cell competition between aneuploid and diploid hESCs occurs in an undifferentiated state and is selectively regulated by Myc independently of other pluripotency transcription factors.

**Figure 6.**
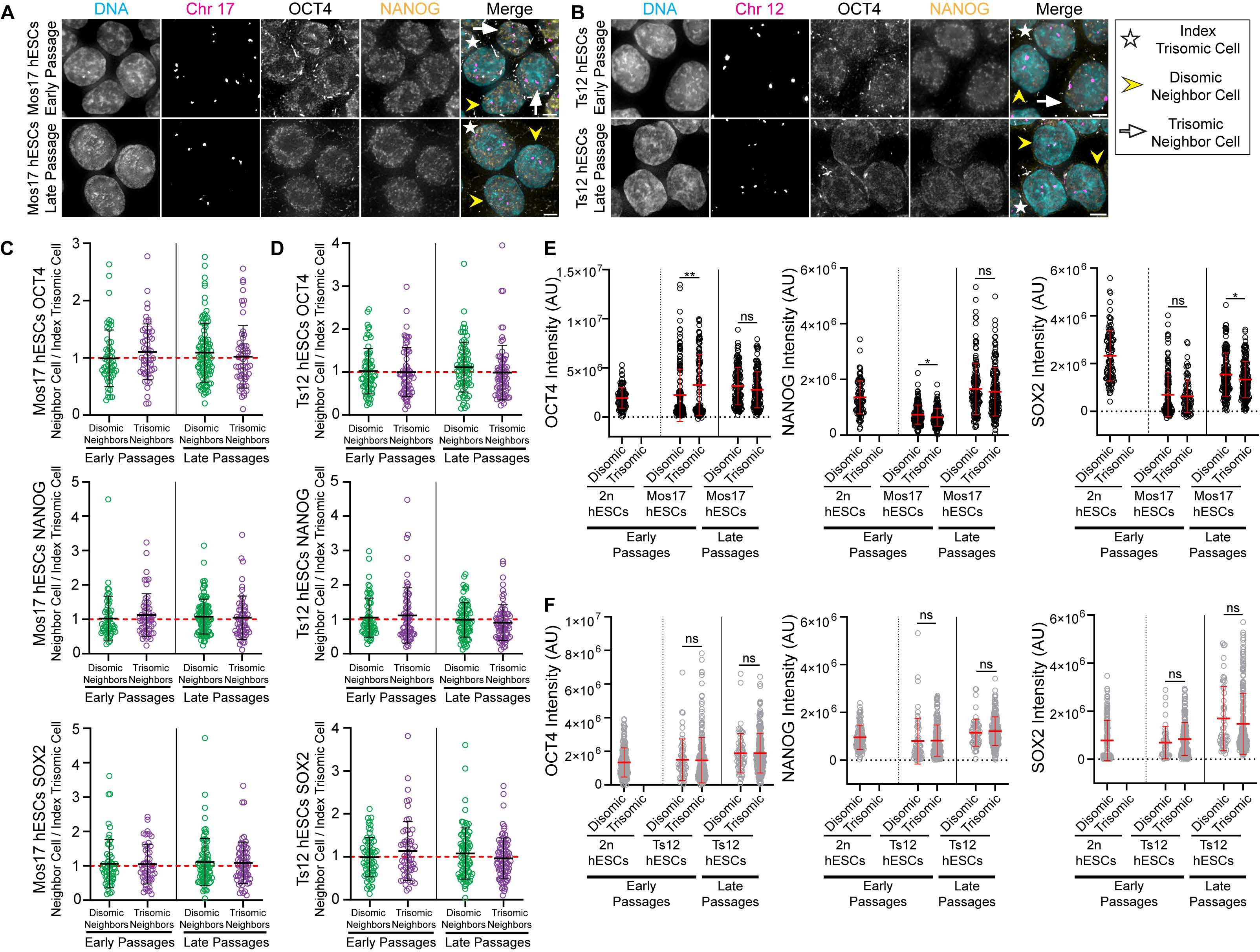
hESC competition occurs in an undifferentiated state. (**A**-**B**) Representative IF-FISH images of Mos17 hESC (**A**) or Ts12 (**B**) neighborhoods comprised of an index trisomic cell (white star), disomic neighbor cells (yellow arrowhead), or trisomic neighbor cells (white arrow) at early (top) or late (bottom) passage. Show is DNA (cyan), chromosome 17 or 12 (magenta), Oct4 (gray), and Nanog (yellow). Scale bar: 5 μm. (**C**-**D**) Quantification of Oct4 (top), Nanog (middle), or Sox2 (bottom) protein levels by IF-FISH. Neighbor cells were normalized to index trisomic cells in early (left) and late (right) passage Mos17 (**C**) or Ts12 (**D**) hESCs. Red dashed line indicates a ratio of 1 and equivalent Oct4, Nanog, or Sox2 levels between index trisomic cells and neighbors. n = 60 index trisomic cells with > 152 neighbors; mean ± SD. (**E**-**F**) Quantification of Oct4 (left), Nanog (middle), or Sox2 (right) protein levels by IF-FISH in randomly selected, non-neighboring 2n and Mos17 hESCs (**E**) or 2n and Ts12 hESCs (**F**) at early (left panels) and late (right panels) passages. n= 300 total cells; mean ± SD. Three independent experiments for both early and late passages (**C**-**F**); ns p > 0.05, *p < 0.05, **p < 0.01 using a one sample t-test compared to hypothetical mean value 1 (**C** and **D**) or a two-tailed Student’s t-test within each cell line for early or late passage (**E** and **F**). See also Figure S6.

## Discussion

Here we show that as a homogeneous population of aneuploid cells, with chromosome gains and losses, hESCs do not elicit a robust cell autonomous response to aneuploidy that inhibits their proliferation. Thus, unlike somatic cells, hESCs are inherently tolerant to aneuploidy. Why hESCs, and presumably induced PSCs, do not respond in a similar manner to chromosome missegregation and aneuploidy as somatic cells is unclear, but our work implicates several possible reasons. We demonstrate that upon induction of aneuploidy, hESCs do not significantly increase p53 absolute protein abundance. Yet, we and others find that p53 absolute levels increase in response to DNA damage leading to cell death^32^. Thus, p53 signaling is functional in hESCs, but aneuploidy does not trigger sufficient p53 activation to cause cell autonomous arrest or apoptosis as occurs in somatic cells^20,23–25^. The lack of a p53 response to aneuploidy in hESCs may in part be due to their low frequency of micronuclei (MN) formation.

Unlike in somatic cells, we find that lagging chromosomes in hPSCs, including both embryonic and induced PSCs, rarely form MN^14^. MN in somatic cells often undergo nuclear envelope rupture incurring DNA damage on the chromosome(s) encapsulated in the micronucleus leading to cell intrinsic responses such as apoptosis or senescence^51^. Accordingly, the reincorporation of lagging chromosomes into the main nucleus in hESCs may prevent MN-associated DNA damage and the activation of downstream anti-proliferative pathways, at least in hESCs that gain an extra chromosome(s). However, DNA damage is not obligatory to increase p53 levels following chromosome missegregation in somatic cells^23,24^, so the low frequency of MN formation is likely to be only one potential mechanism responsible for the tolerance of hESCs to aneuploidy. Moreover, we find that, compared to somatic cells, p21 protein levels downstream of p53 are repressed in hESCs in response to both DNA damage^31,33,34^ and aneuploidy likely contributing to their aneuploidy tolerance. Interestingly, increased expression of p53 or p21 above basal levels both promotes differentiation of hPSCs^52,53^. Thus, the transcriptional networks required to maintain a pluripotent state may coincidentally render hESCs permissive to aneuploidy.

An inherent tolerance to aneuploidy combined with hESCs exhibiting a persistently high mitotic error rate^14^ poses a conundrum in how hESCs maintain stable diploid populations for generations. Mechanistically, our data demonstrates that Myc- and p53-driven cell competition eliminates less fit aneuploid hESCs to preserve the genome integrity of the population (presumably a similar mechanism acts in induced PSCs) (Figure 7A). Although we show that this occurs for hESCs that gain chromosome 17 copies, Myc- and p53-mediated cell competition likely occurs independent of specific chromosome imbalances because we find that chromosome missegregation is random in hESCs, yet they maintain diploid populations^14^.

**Figure 7.**
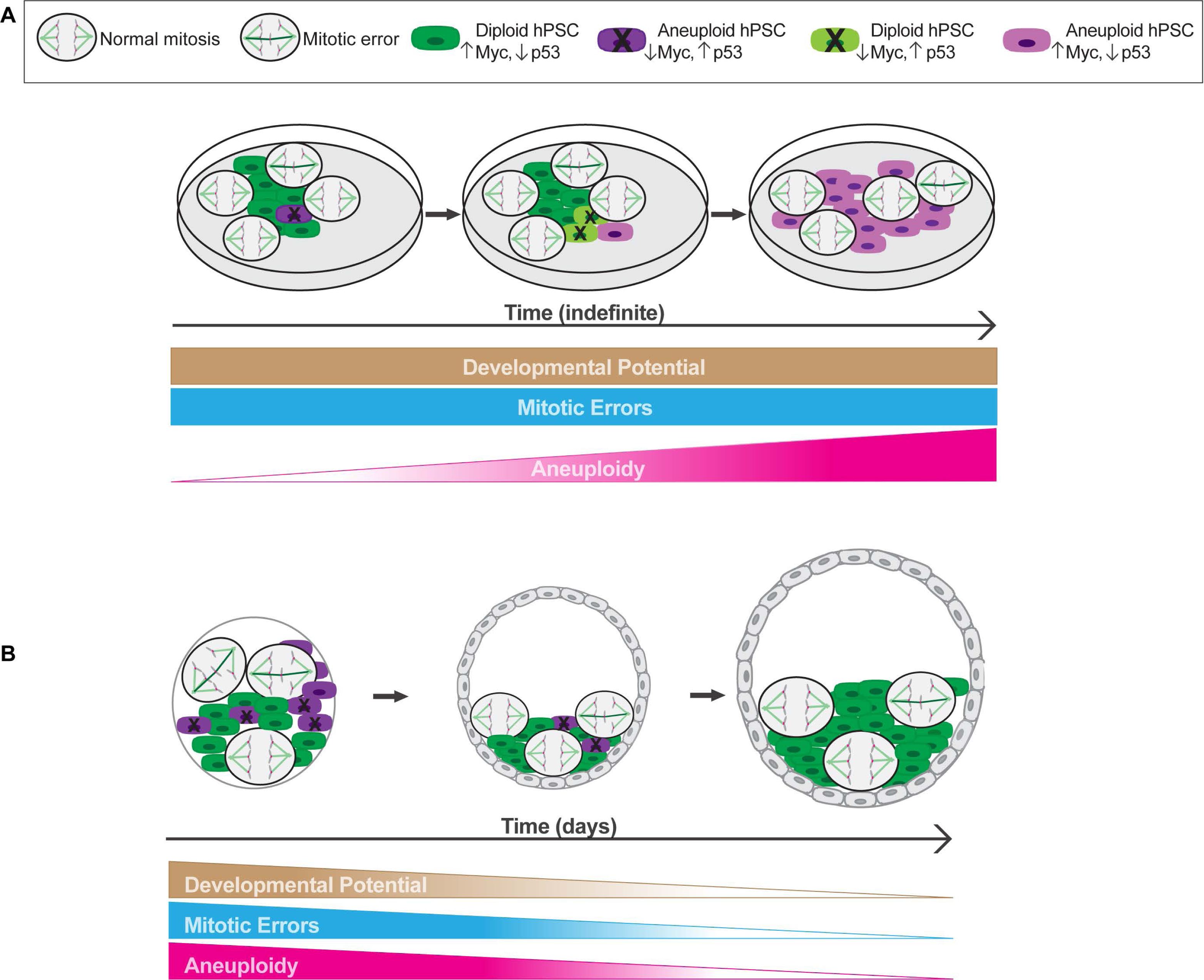
Models to preserve genome stability in hPSCs and to establish diploid preimplantation embryos. (**A**) HPSCs grow in culture indefinitely and maintain a pluripotent state that confers a persistent and high rate of mitotic errors, particularly lagging chromosomes, generating aneuploid progeny. Yet, genome integrity is preserved because less fit aneuploid hPSCs with decreased Myc and increased p53 abundance relative to more fit neighboring diploid hPSCs are outcompeted. However, during prolonged passaging hPSCs, acquire mutations that cause a switch in fitness resulting in aneuploid hPSCs with increased Myc and decreased p53 relative abundance gaining a selective advantage and overtaking the population. (**B**) Preimplantation embryonic cells also have a high rate of mitotic errors, including lagging chromosomes, that generate aneuploid progeny. We propose that, like hPSCs, Myc- and p53-driven cell competition eliminates aneuploid embryonic cells. This, coupled with a declining mitotic error rate as developmental potential becomes restricted, decreases the frequency of aneuploid embryonic cells to achieve genome stability and establish diploid embryos as development progresses. Models adapted from Deng et al, 2023^14^.

Furthermore, our findings suggest that premature differentiation is not responsible for the initial fitness disadvantage conferred by an aneuploid genome. Alternatively, in non-transformed somatic cells, aneuploidy elicits stress responses, including proteotoxic and metabolic stress, independent of specific chromosome imbalances^21,54^, and these stress pathways decrease cell fitness and promote cell competition in other organisms^55–57^. Thus, we speculate that aneuploidy induced stress responses could contribute to relative protein abundance differences between neighboring aneuploid and diploid hESCs to promote cell competition, particularly since Myc and p53 play key roles in regulating cellular metabolism and protein homeostasis^58,59^.

Additionally, cell competition is primarily dependent on short-range interactions between neighbors^36,42,47^, and our data suggest this is the same for Myc- and p53-mediated competitive interactions between aneuploid and diploid hESCs. But it is unknown how relative differences in cell fitness are communicated between neighbors to affect their fate.

We also find that the fitness costs of an aneuploid genome can be overcome during extended passaging leading to aneuploid hESCs with relatively higher Myc and lower p53 levels gaining a competitive advantage over diploid hESCs (Figure 7A). Previous work demonstrates that aneuploid hPSCs with a competitive advantage eliminate diploid hPSCs due to YAP mediated mechanical competition^3^, but it is unknown if YAP acts in the same pathway as Myc and p53 or if these are independent mechanisms. Regardless, the switch in fitness for aneuploid relative to diploid hESCs most likely requires the acquisition of additional genetic changes (assuming routine and consistent dissociation and culturing). Our data suggests that these changes are likely focal copy number variations or point mutations because no other clonal whole chromosome abnormalities arose. We favor a model where aneuploid hESCs gain an additional mutation(s) to improve their relative cell fitness because hESCs frequently acquire cancer-related mutations^29,30^ and aneuploidy is also a common phenotype of cancers^60^. A requirement for multiple genetic changes also explains why aneuploid hESCs typically do not overtake a population until after extended passaging^1,4^. Furthermore, assuming the spontaneous mutation rate is constant during routine culturing, this may account for why we observe a fairly consistent time frame for when trisomic Mos17 hESCs gain a competitive advantage. This model also predicts that once there is a sufficient proportion of aneuploid hESCs with a fitness advantage, the trait is fixed in the population, because although ongoing chromosome missegregation may revert aneuploid hESCs back to a diploid state, the diploid hESCs generated will be relatively less fit and outcompeted. In support, we do not observe Ts12 hESCs returning to a predominantly diploid population, at least on the time scale of our analysis. Interestingly, however, we find that overcoming the fitness costs of aneuploidy does not automatically promote genomic instability. Rather, there must be additional selection pressures that favor the proliferation of hESCs with certain aneuploidies explaining the preferential occurrence of karyotypes in culture-adapted aneuploid hESCs^1^. Yet, genetic variation between hESC lines must also play a role in this process because the CSES7 2n hESCs remained a stable diploid population despite being cultured for an equivalent time. Collectively, these results pose new considerations for how to define acceptable thresholds of stem cell genomic integrity in regenerative medicine therapies.

Our results also have important implications for human preimplantation development where >70% of embryos are aneuploid and mosaic diploid and aneuploid embryos are common due to the prevalence of mitotic chromosome missegregation^5–8^. Based on our findings, we propose that Myc- and p53-driven cell competition depletes aneuploid cells in mosaic embryos during preimplantation development (Figure 7B). In support, the most significantly down- regulated pathway in aneuploid vs. diploid human preimplantation embryonic cells is Myc targets indicative of relative differences in Myc protein abundance^6^ and consistent with our findings in hESCs. We note that *in vivo* the time scale for cell competition-based elimination would have to be accelerated compared to our *in vitro* results given that preimplantation development happens over the course of days. Moreover, the propensity for chromosomes to missegregate is mostly comparable^5,8^ and Myc- and p53-mediated cell competition likely occurs independent of specific chromosome imbalances. Furthermore, there is no preferential lineage allocation of aneuploid embryonic cells to either the trophectoderm (TE) or pluripotent inner cell mass (ICM)^6,8,61^. Since hESCs more closely resemble pluripotent ICM cells an open question is whether cell competition occurs between aneuploid and diploid TE cells. Subsequently as development progresses and developmental potential becomes restricted, mitotic error rates decrease^14^ thus reducing the generation of aneuploid cells coupled with a switch to cell autonomous mechanisms that eliminate aneuploid cells (Figure 7B). One critical parameter in this model is the ratio of diploid to aneuploid cells during preimplantation development. There must be a sufficient proportion of more fit diploid embryonic cells in contact with less fit aneuploid cells to eliminate them or at least to support developmental progression until cell autonomous mechanisms act. A disproportionate number of aneuploid embryonic cells may explain why not all transferred IVF mosaic embryos result in healthy live births with diploid karyotypes^18^. In contrast, hPSCs indefinitely maintain a pluripotent state resulting in both a persistently high mitotic error rate^14^ and the continuous generation of aneuploid cells. Thus, increasing the likelihood that during extended passaging aneuploid hPSCs acquire additional mutations that overcome the fitness barriers of an aneuploid genome (Figure 7A). In conclusion, we find genome integrity with respect to chromosome copy numbers is controlled by distinct mechanisms dependent on developmental state.

## Acknowledgements

We thank Nissim Benvenisty for providing cell lines, Yina Huang for sharing instrumentation, and Aaron McKenna for sharing plasmids. Also, we thank Duane Compton, Compton lab members, Aaron McKenna, and Erik Griffin for their helpful discussions. This work was supported by National Institute of Health R01HD101436 to KMG.

## Author contributions

Conceptualization-KMG; Methodology-KMG, CD, AY; Validation-KMG, CD, AY; Formal Analysis-KMG, CD, AY; Investigation-CD, AY; Resources-KMG; Writing-Original Draft-KMG, AY; Writing-Review and Editing-KMG, CD, AY; Visualization-KMG, CD, AY; Supervision-KMG; Funding Acquisition-KMG.

## Supplemental information

Document S1: Figures S1–S6.

## Methods

### Cell lines and cell culture

RPE-1 H2B-GFP (XX) cells were grown in DMEM supplemented with 10% fetal calf serum (FCS), 50 U/mL penicillin, 50 μg/mL streptomycin, 250 μg/L Amphotericin B, and 10 mM HEPES and passaged using 0.05% trypsin. H1/WA01 (XY) and H1 H2B-GFP (XY) hESCs were grown in mTeSR1 on hESC qualified Matrigel. CSES7 (XX), CSES8 (XX), and CSES22 (XX) hESCs were grown in mTeSR Plus on hESC qualified Geltrex™. For routine passaging, H1, H1 H2B-GFP, CSES7, CSES8, and CSES22 hESCs were dissociated using versene according to the WiCell protocol SOP-SH-002 (version H). All cell lines were validated as mycoplasma free and grown at 37°C in a humidified atmosphere with 5% CO_2_. G-banded karyotyping and short tandem repeat analysis were performed by WiCell Research Institute.

We generated RPE-1 cells and H1 hESCs that stably express GFP-H2B by co-transfecting cells with the PiggyBac-EF1α-H2B-GFP-mCherry-Cdt1 with G418 or puromycin resistance cassettes and PiggyBac transposase plasmids. To construct the plasmids, H2B-GFP was linearized using NheI and NotI restriction enzymes. DNA inserts EF1α and H2B-GFP were PCR amplified and cloned into PB-pCMV-MCS-EF1α-Puro using NheI and SpeI restriction enzymes creating the intermediate plasmid PB-EF1α-H2B-GFP. Subsequently, to construct PB- EF1α-H2B-GFP-mCherry-Cdt1 with puromycin or G418 resistance, the PB-EF1α-H2B-GFP vector was linearized using the NotI restriction enzyme. DNA inserts mCherry-Cdt1, IRES, and NeoR with overlapping ends were PCR amplified from pEN435-pCAGGS-TagBFP-hGeminin- 2A-mCherry-hCdt1-rbgpA-Frt-PGK-EM7-PuroR-bpA-Frt Tigre targeting, BII-ChBtW-iresFUCCI, and pTet-ON plasmids, respectively. The DNA products were cloned into the linearized PB- EF1α-H2B-GFP vector using Gibson assembly.

RPE-1 cells were transfected using Lipofectamine 2000 following the manufacturer’s instructions and H1 hESCs were transfected using nucleofection following the manufacturer’s instructions and the protocol from Zhong et al, 2020^62^. RPE-1 cells stably expressing H2B-GFP were selected using 1 mg/mL G418 and H1 hESCs stably expressing H2B-GFP were selected using 0.3 μg/mL puromycin. Note, both H1 H2B-GFP hESCs and RPE-1 H2B-GFP cells also express a G1 reporter, but we did not monitor G1 phase in our experiments.

### Immunofluorescence (IF), fluorescence *in situ* hybridization (FISH), and immunofluorescence-FISH (IF-FISH)

For IF, FISH, or IF-FISH experiments, H1, H1 H2B-GFP, CSES7, CSES8, and CSES22 hESCs were plated as aggregates on Matrigel-coated 18 mm glass coverslips in 12-well cell culture plates. RPE-1 H2B-GFP cells were plated on standard 18 mm glass coverslips in 12-well cell culture plates. IF experiments were performed as previously described by our lab^14^. Primary antibodies were diluted in TBS + 2% BSA + 0.1% Triton X-100 at 5 μg/mL rabbit anti-OCT4, 1:200 mouse anti-NANOG, 1:1600 rabbit anti-MYC, 1:1000 mouse anti-p53, 10 μg/mL mouse anti-SOX2, or 1:400 rabbit anti-p21.

For FISH experiments, upon completion of drug treatment or reaching optimal confluency, cells were fixed with 3.5% PFA for 5 mins at room temperature, permeabilized with TBS + 0.1% Triton X-100 for 2 × 5 mins and washed with TBS for 3 × 5 mins. Subsequently, FISH was performed as previously described by our lab^63^.

For IF-FISH experiments, the standard IF protocol as previously described was perfomed^14^, with the exception that secondary antibodies were diluted in TBS + 2% BSA + 0.1% Triton X-100 without DAPI. Subsequently, cells were washed with TBS + 2% BSA + 0.1% Triton X-100 for 3 × 5 mins and TBS buffer for 2 x 5 mins sequentially followed by performing our standard FISH protocol as previously described^63^.

### Microscopy for IF, FISH, and IF-FISH

Images were acquired with a Hamamatsu ORCA-Fusion Gen III sCMOS camera on a Nikon Eclipse Ti2E inverted microscope using a Nikon Plan Apo Lambda 60×, 1.4 numerical aperture oil immersion objective. Image series in the Z axis were obtained using 0.5 µm steps for a total of 9 optical slices. For quantification of proteins, all images were acquired with the same acquisition parameters and exposure times. For quantification of chromosome copy number only, exposure times were adjusted based on the probe signal intensity.

Image deconvolution and contrast enhancement were performed using NIS Batch Deconvolution (Nikon), NIS Elements (Nikon), and ImageJ (NIH). Images shown are maximum intensity projections of selected Z-planes.

### Time-lapse live-cell fluorescence imaging

Live-cell imaging was performed at 37°C in a humidified environment with 5% CO2 (Tokai Hit Stage-top Incubation System) using a Nikon Plan Apo Lambda 60×, 1.4 numerical aperture oil immersion objective, a Hamamatsu ORCA-Fusion Gen III sCMOS camera on a Nikon Eclipse Ti2E inverted microscope with binning set to 2×2. Image series in the *Z* axis were obtained using 1 µm steps for a total of 7 optical slices.

H1 H2B-GFP hESCs were plated in mTeSR1 on the 35 mm glass bottom dishes coated with Matrigel. RPE-1 H2B-GFP cells were plated in DMEM on the 35 mm glass bottom dishes. Cells were incubated for 1-2 days at 37°C in a humidified atmosphere with 5% CO2 prior to live-cell imaging. Subsequently, cells were washed with phenol-free mTeSR1 (H1 H2B-GFP) or FluoroBrite™ DMEM (RPE-1 H2B-GFP) three times and then cultured in phenol-free medium supplemented with 0.1% DMSO, 75 nM, or 100 nM reversine during live-cell imaging. Cells were imaged for 6-7 hours with 2 min time intervals. We optimized these experiments using the lowest exposure and intensity settings permissible to visualize errors while minimizing artifacts due to phototoxicity.

Image acquisitions and analyses were performed using NIS Elements (Nikon) and ImageJ (NIH). Representative images from live-cell experiments are maximum intensity projections of all Z-planes. Cells undergoing mitosis were tracked from nuclear envelope breakdown (NEB) to anaphase onset (AO), during which prometaphase (NEB to metaphase), metaphase (metaphase to AO) or total mitotic (NEB to AO) durations were recorded. In combination with mitotic duration, anaphase errors were observed and scored. Criteria for scoring chromosome segregation errors was previously described^14^.

### Quantification of chromosome copy number

To determine chromosome copy number, sum intensity projections were compiled from Z-stack images using ImageJ (NIH). Nuclei were randomly selected per field of view based upon the DAPI signal. Chromosome copy number was determined based on the number of fluorescent puncta counted per individual nuclei.

### Quantification of protein expression in non-neighboring cells

To determine protein expression levels, sum intensity projections were compiled from *Z*- stack images using ImageJ (NIH). Nuclei were randomly selected per field of view based upon the DAPI signal. An elliptical region of interest (ROI) was drawn to encompass the entire nucleus and then a slightly larger elliptical ROI was drawn to encompass both the nucleus and the adjacent background. The mean background intensity was calculated based on the in-between region of the two ROIs and subtracted from the integrated intensity of the ROI that encompassed the nucleus. It is important to note that due to background correction, the background-subtracted intensity of the nuclear ROI may fall below zero for proteins with low expression levels or if there was pronounced non-nuclear background signal. Chromosome copy numbers in selected cells were determined as described above.

### Quantification of protein expression in neighboring cells

To determine a cell “neighborhood”, an index trisomic chromosome 17 or 12 cell was chosen as described above and if it was directly neighbored by at least one disomic cell. Once an index trisomic cell was selected, every neighboring cell, regardless of chromosome 17 or 12 copy number, was included in the neighbor analysis to quantify relative protein expression levels. The number of cells included in each neighborhood varied with a minimum of at least two neighboring cells analyzed per neighborhood. Quantification of protein expression of all cells in a neighborhood was performed as described above. Protein expression levels for each neighboring cell were then normalized to the protein expression level of the index trisomic cell. This quantification was repeated for each cell neighborhood analyzed with no cells overlapping between neighborhoods.

### Drug treatments

Cells were treated with 0.1% DMSO, reversine, or doxorubicin at the concentrations and times specified. For growth curve experiments, cells were cultured in mTeSR1 (H1 and H2B- GFP hESCs) or DMEM (RPE-1 H2B-GFP cells) supplemented with DMSO or 100 nM reversine for 96 hours. Fresh supplemented media was added daily for all cell lines. After 96 hours the DMSO or reversine supplemented media was washed out and cells were split into a new 6-well plate and cultured with fresh media in the absence of DMSO or reversine for an additional 96 hours. Fresh media was added daily for all cell lines.

For all doxorubicin experiments, cells were cultured in mTeSR1 (H1 and H1 H2B-GFP hESCs) or mTeSR Plus (CSES7, CSES8, CSES22 hESCs) supplemented with 1 μM doxorubicin for 4 hours followed by washout with fresh media before IF. For the cell viability assay, cells were cultured in doxorubicin for 24h.

### Statistical analysis

GraphPad Prism was used for all statistics. The statistical tests used, corresponding n values, error bar measurements, and p values are in the figure legends. No outliers were excluded.

### Reagents

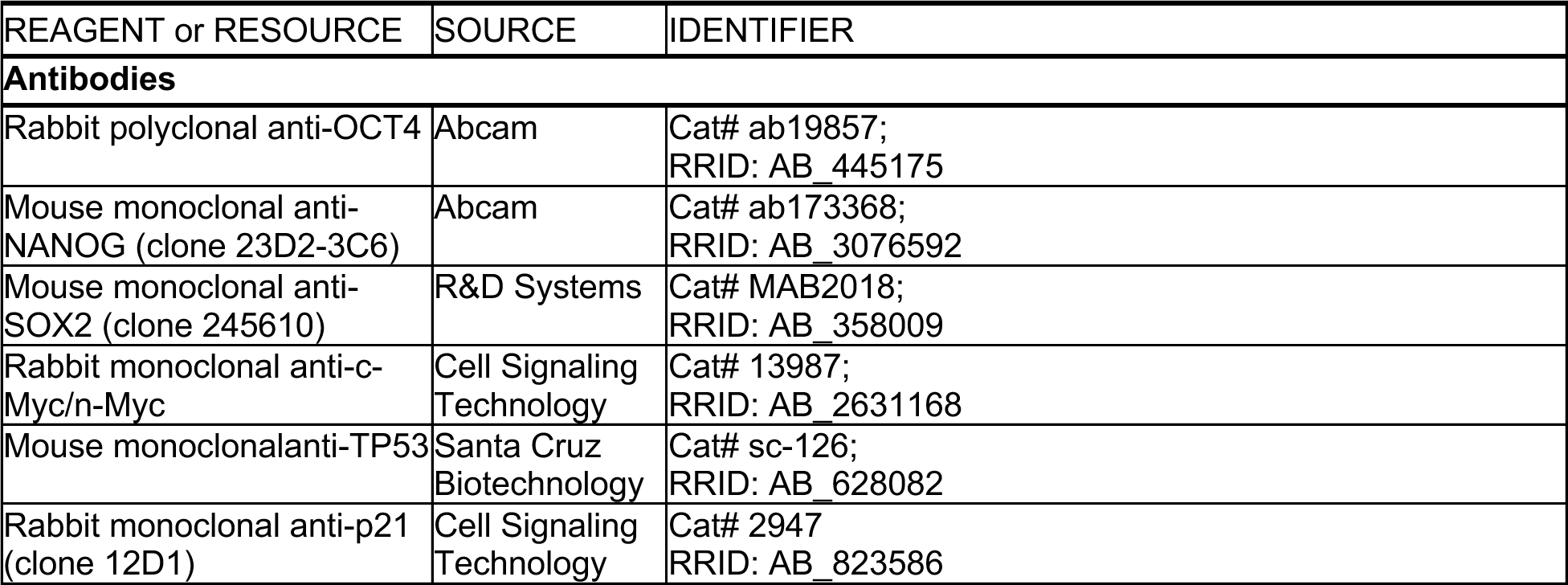
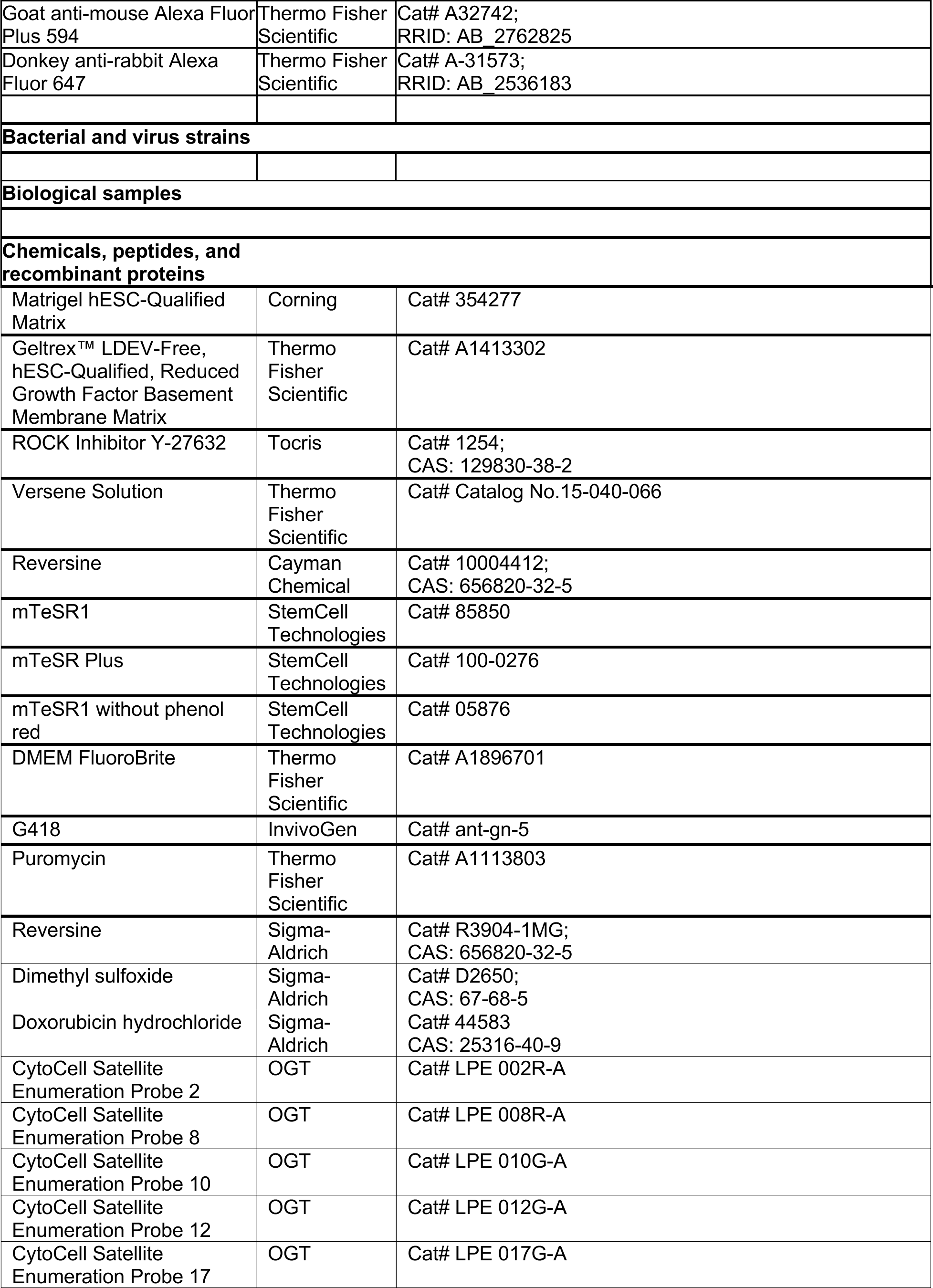
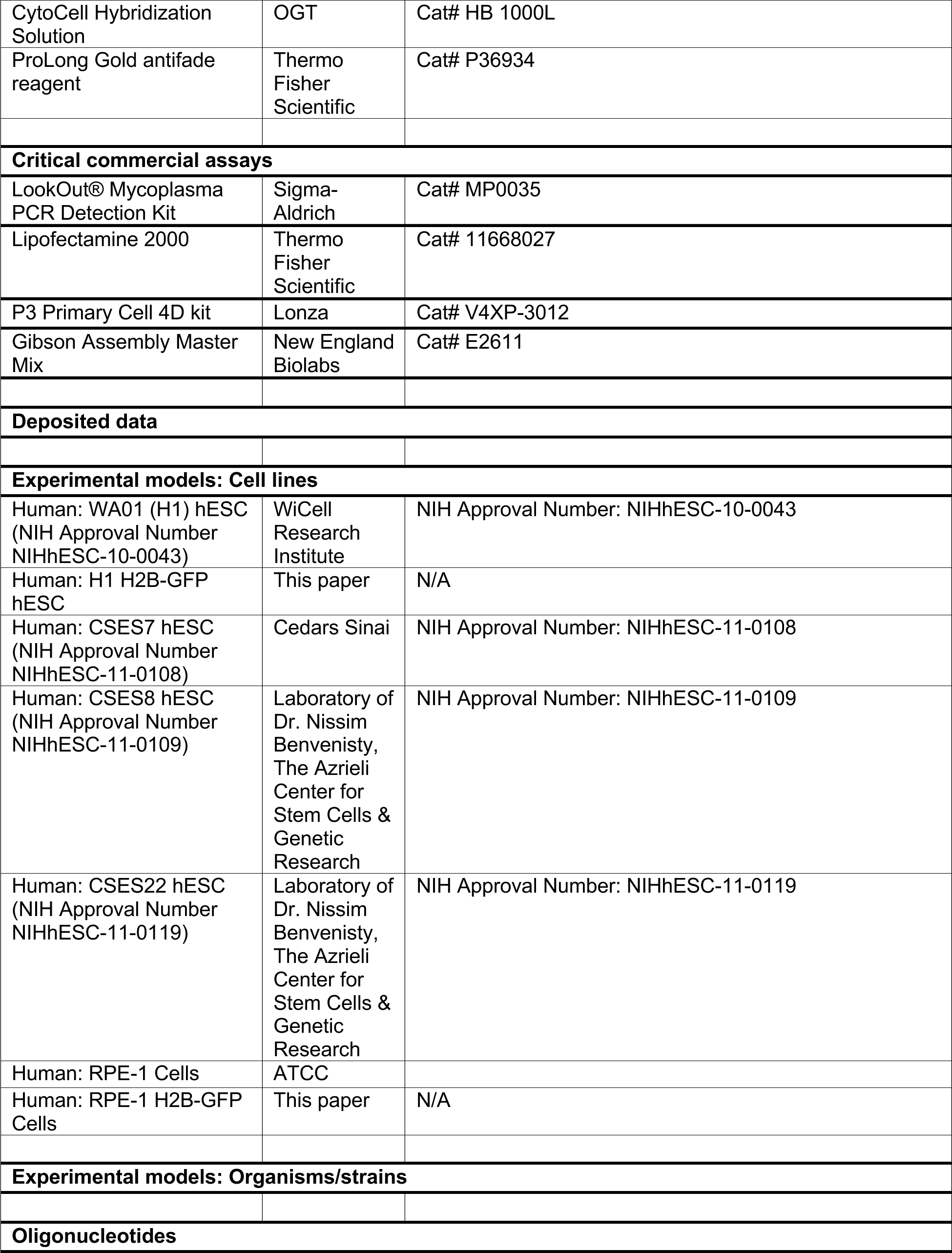
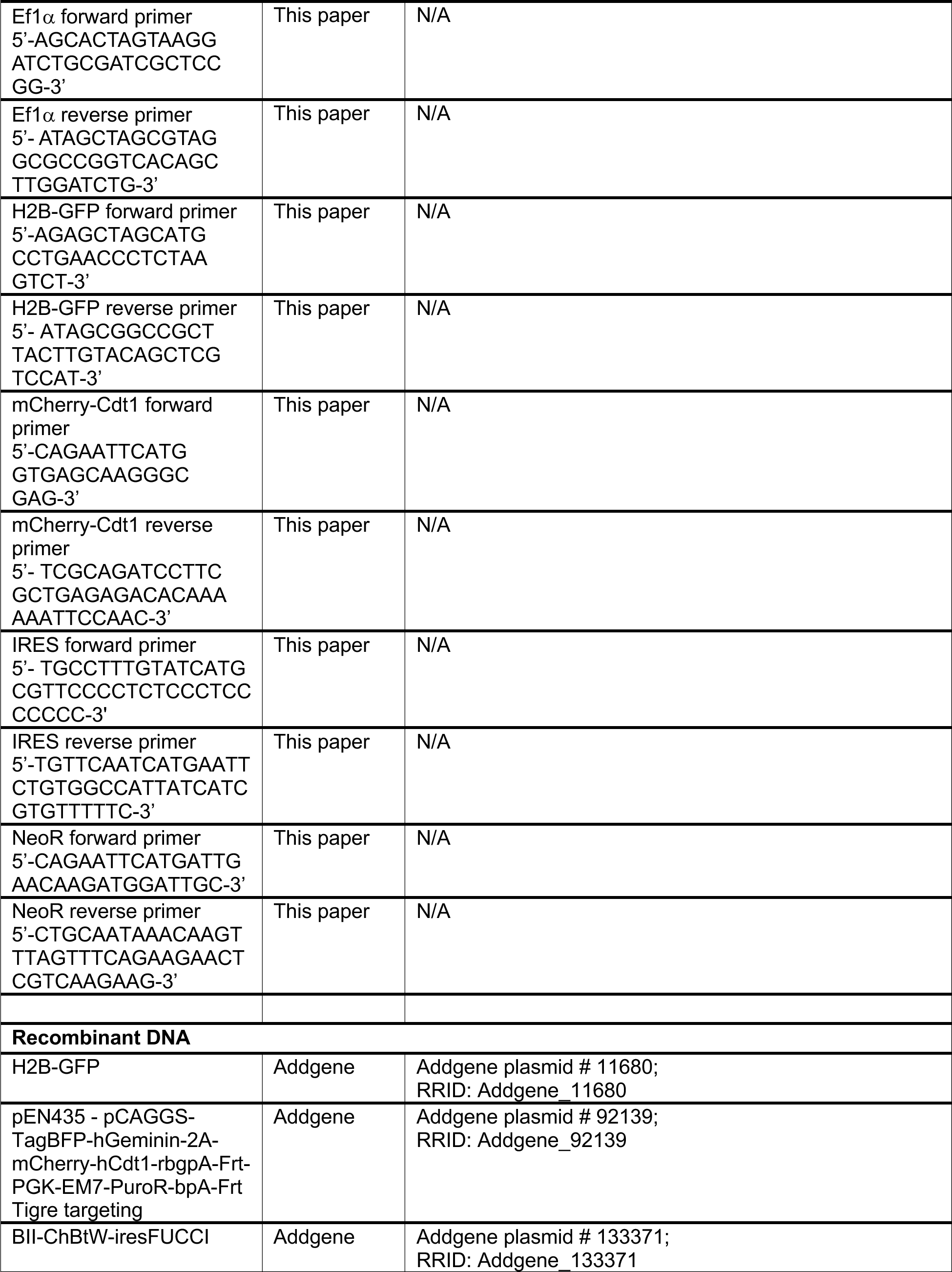
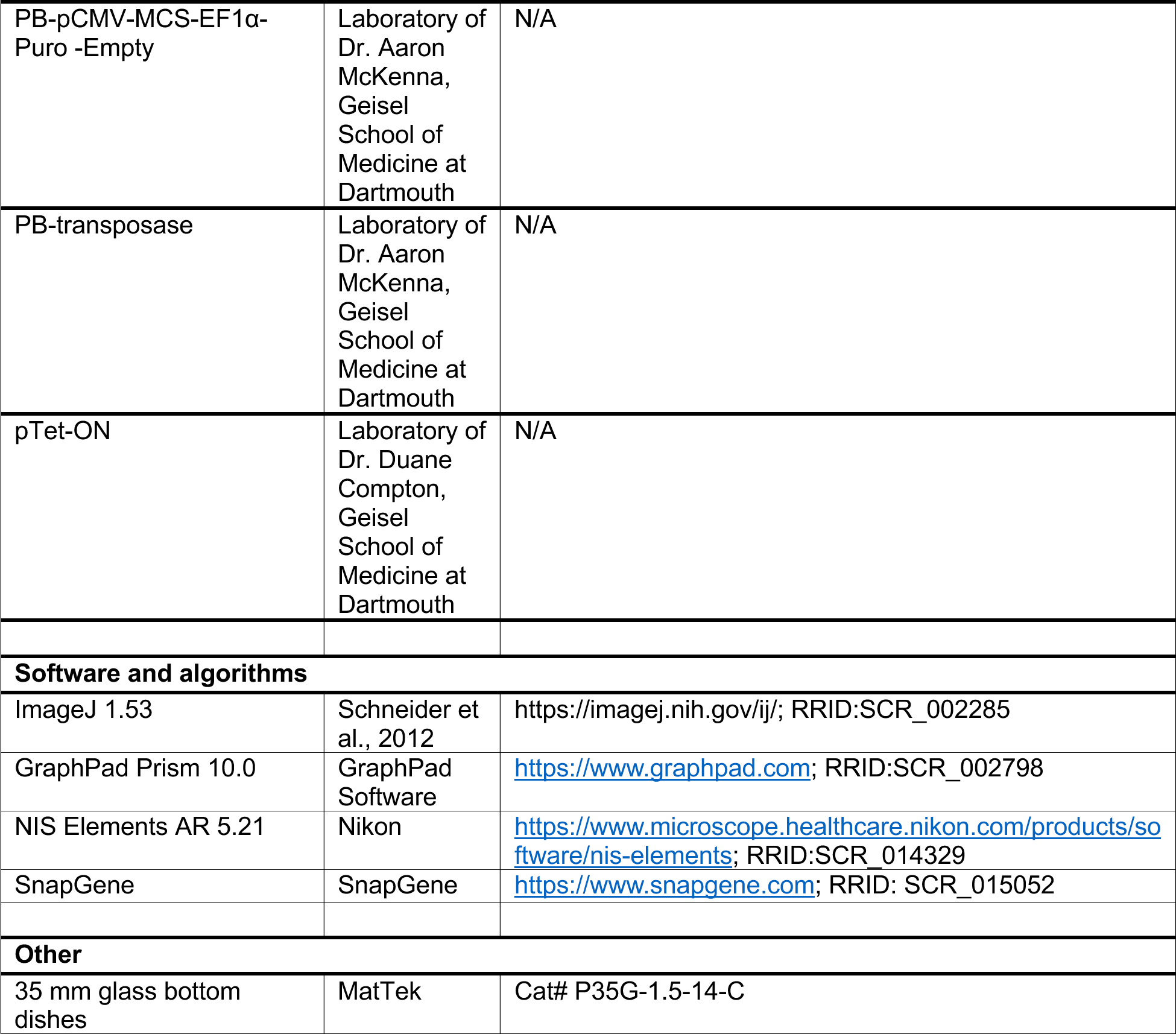

