## Supplemental Figures for "Cell Competition Eliminates Aneuploid Human Pluripotent Stem Cells"

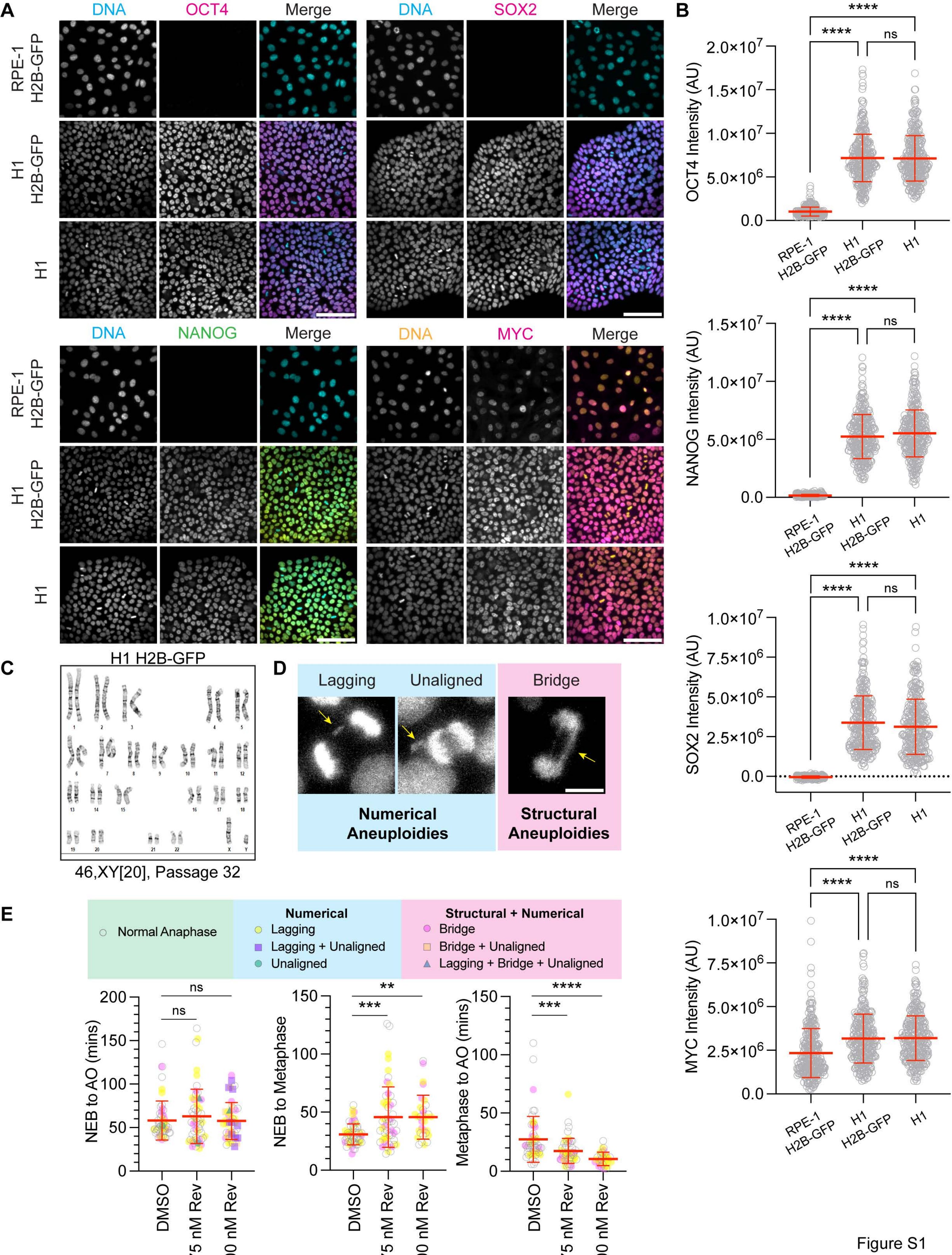
A
DNA
OCT4
Merge
DNA
SOX2
Merge
RPE-1 H2B-GFP
H1 H2B-GFP
H1
DNA
NANOG
Merge
DNA
MYC
Merge
B
OCT4 Intensity (AU)
NANOG Intensity (AU)
SOX2 Intensity (AU)
MYC Intensity (AU)
****
ns
C
H1 H2B-GFP
46,XY[20], Passage 32
D
Lagging
Unaligned
Bridge
Numerical Aneuploidies
Structural Aneuploidies
E
Normal Anaphase
Numerical
Lagging
Lagging + Unaligned
Unaligned
Structural + Numerical
Bridge
Bridge + Unaligned
Lagging + Bridge + Unaligned
NEB to AO (mins)
NEB to Metaphase
Metaphase to AO (mins)
DMSO
75 nM Rev
100 nM Rev
***
**

Figure S1

**Figure S1. Pluripotency transcription factor expression, karyotype analysis, anaphase error rates, and mitotic durations in H1 hESCs, H1 H2B-GFP hESCs, or RPE-1 H2B-GFP cells. Related to Figure 1.**

(**A**) Representative IF images of somatic RPE-1 H2B-GFP cells, H1 H2B-GFP hESCs, and H1 hESCs that show DNA (cyan or yellow), Oct4 (magenta), Nanog (green), Sox2 (magenta), or Myc (magenta). Scale bars: 100 µm. (**B**) Quantification of Oct4, Nanog, Sox2, or Myc protein levels by IF in somatic RPE-1 H2B-GFP cells, H1 H2B-GFP hESCs, and H1 hESCs. $n = 300$ cells; mean ± SD. (**C**) G-banding karyotype analysis of twenty H1 H2B-GFP hESCs showing no clonal abnormalities. (**D**) Representative IF images of mitotic errors that cause numerical or structural aneuploidies from time-lapse live-cell microscopy of H1 H2B-GFP hESCs treated with 100 nM reversine. Scale bar: 10 µm. (**E**) Mitotic duration (left), prometaphase duration (middle) and metaphase duration (right) of H1 H2B-GFP hESCs treated with DMSO, 75 nM, or 100 nM reversine. NEB, nuclear envelope breakdown; AO, anaphase onset; $n > 45$ cells; mean ± SD. Three independent experiments (**B** and **E**); ns $p > 0.05$, $^{**}p < 0.01$, $^{***}p < 0.001$, and $^{****}p < 0.0001$ one-way ANOVA and Tukey’s multiple comparisons test (**B**) or a one-way ANOVA and Dunnett’s multiple comparisons test (**E**).

A

| | Mitotic duration (mins) | Prometaphase duration (mins) | Metaphase duration (mins) | Numerical Errors (%) | Structural Errors (%) |
|---|---|---|---|---|---|
| RPE-1 H2B-GFP DMSO *Deng et al. 2023.* | 22.4 | 12.2 | 10.2 | 7% | 4% |
| RPE-1 H2B-GFP 100 nM Rev | 23.4 | 17.3 | 6.1 | 40.5% | 21.4% |

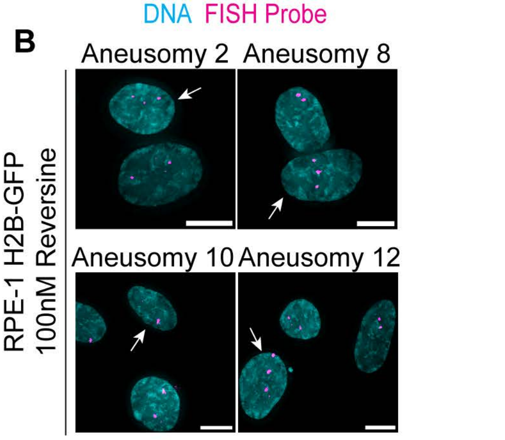


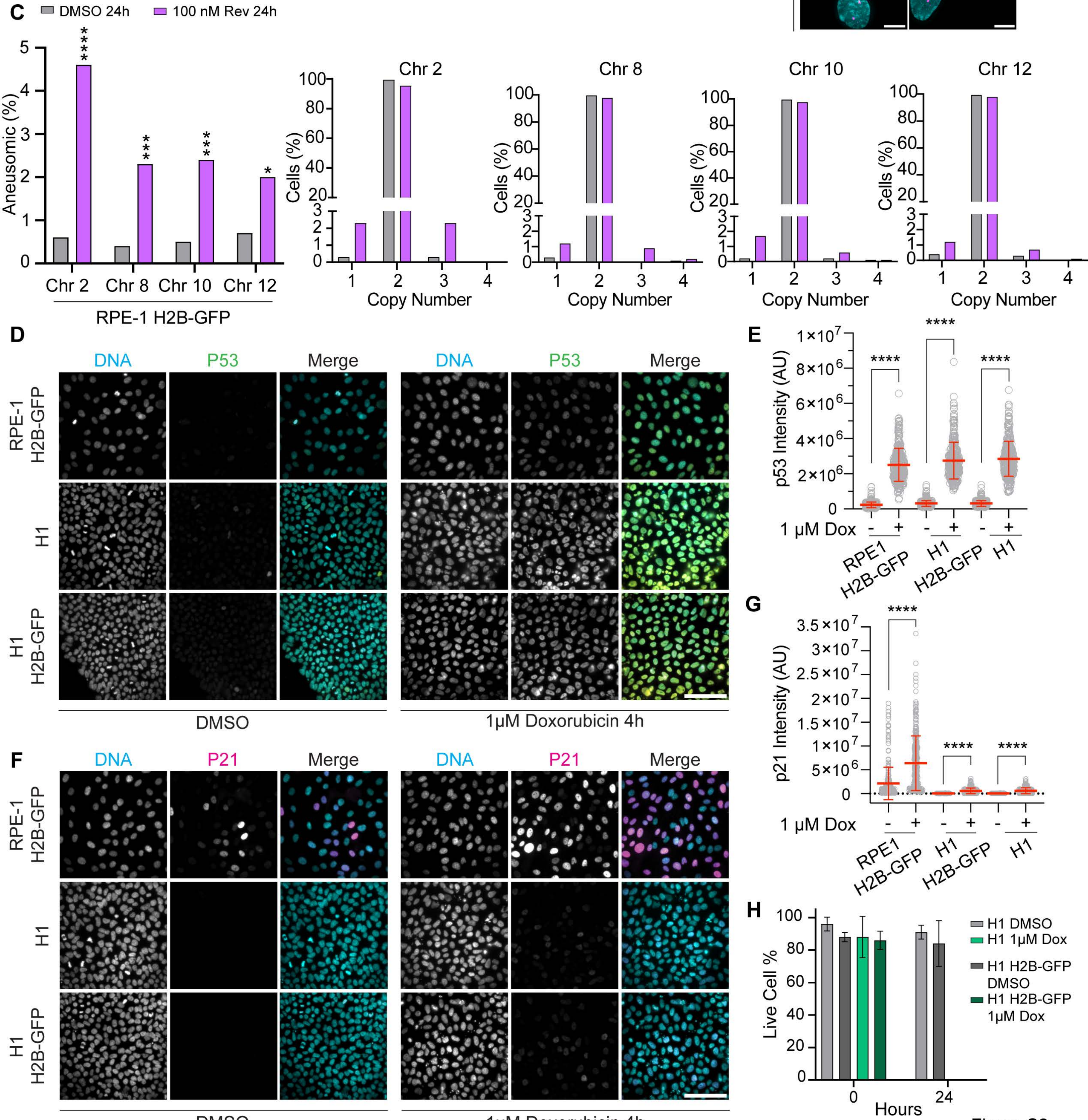


Figure S2

**Figure S2. Mitotic durations, anaphase error rates, aneusomic rates, and p53 and p21 levels in somatic RPE-1 H2B-GFP cells, H1 hESCs, or H1 H2B-GFP hESCs. Related to Figure 2.** (**A**) Mean mitotic duration, prometaphase duration, or metaphase duration, and mitotic error rates of RPE-1 H2B-GFP cells treated with DMSO or 100 nM reversine (Rev). (**B-C**) The copy numbers of chromosomes 2, 8, 10, or 12 in RPE-1 H2B-GFP cells treated with DMSO or 100 nM reversine for 24h determined using FISH. Interphase cells were scored for copy numbers. (**B**) Representative FISH images show aneuploid trisomic or monosomic (white arrows) cells. Scale bars: 10 μm. (**C**) Aneusomic percent, including gains and losses, and copy number distribution of chromosomes 2, 8, 10, or 12. For each chromosome, the aneusomic percent of their respective DMSO and reversine treated cells were compared. n = 1000 cells pooled per FISH probe. (**D**-**E**) Representative IF images (**D**) and quantification of p53 levels (**E**) in somatic RPE-1 H2B-GFP cells, H1 H2B-GFP hESCs, and H1 hESCs treated with DMSO or 1 μM doxorubicin for 4h. Shown is DNA (cyan) and p53 (green). n = 300 cells; mean ± SD. (**F**-**G**) Representative IF images (**F**) and quantification of p21 levels (**G**) in somatic RPE-1 H2B-GFP cells, H1 H2B-GFP hESCs, and H1 hESCs treated with DMSO or 1 μM doxorubicin for 4h. Shown is DNA (cyan) and p21 (magenta). n = 300 cells; mean ± SD. (**H**) Percent of live H1 or H1 H2B-GFP hESCs after DMSO or 1 μM doxorubicin for 24h. Mean ± SD. Two (**C** and **H**) or three (**E** and **G**) independent experiments; *p < 0.05, ***p < 0.001, ****p < 0.0001 using a two-tailed Fisher's exact test (**C**) or two-tailed Student's t-test within each cell line (**E** and **G**).

**A**

| STR Loci | Peaks |
|---|---|
| FGA | 23, 24 |
| TPOX | 8, 8 |
| D8S1179 | 12, 13 |
| vWA | 15, 17 |
| Amelogenin | X, X |
| Penta_D | 9, 11 |
| CSF1PO | 10, 11 |
| D16S539 | 10, 13 |
| D7S820 | 10, 11 |
| D13S317 | 10,11 |
| D5S818 | 12, 13 |
| Penta_E | 12, 16 |
| D18S51 | 13, 15 |
| D21S11 | 31.2, 33.2 |
| TH01 | 7, 7 |
| D3S1358 | 16, 17 |

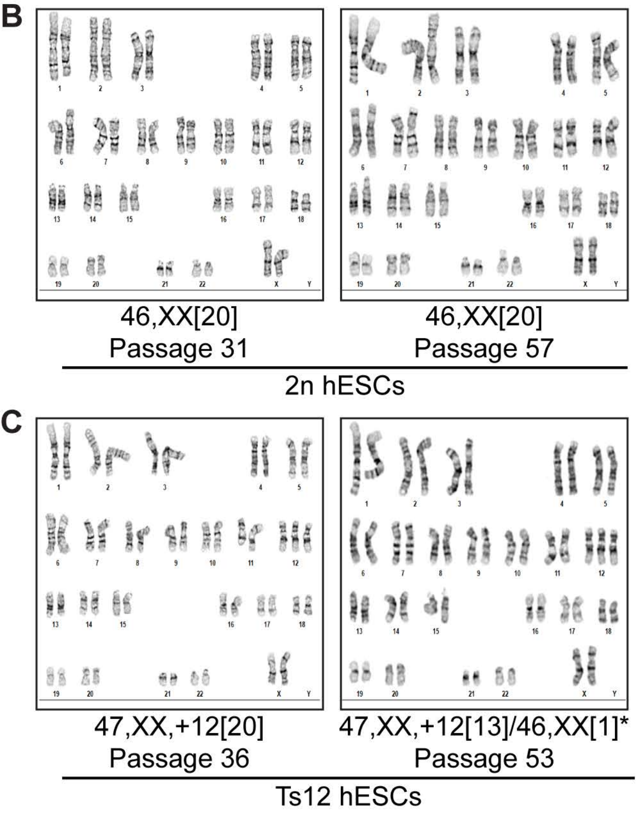


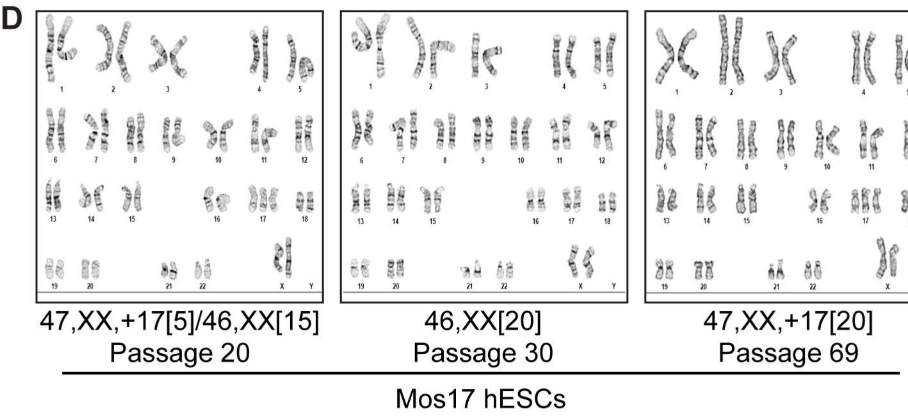


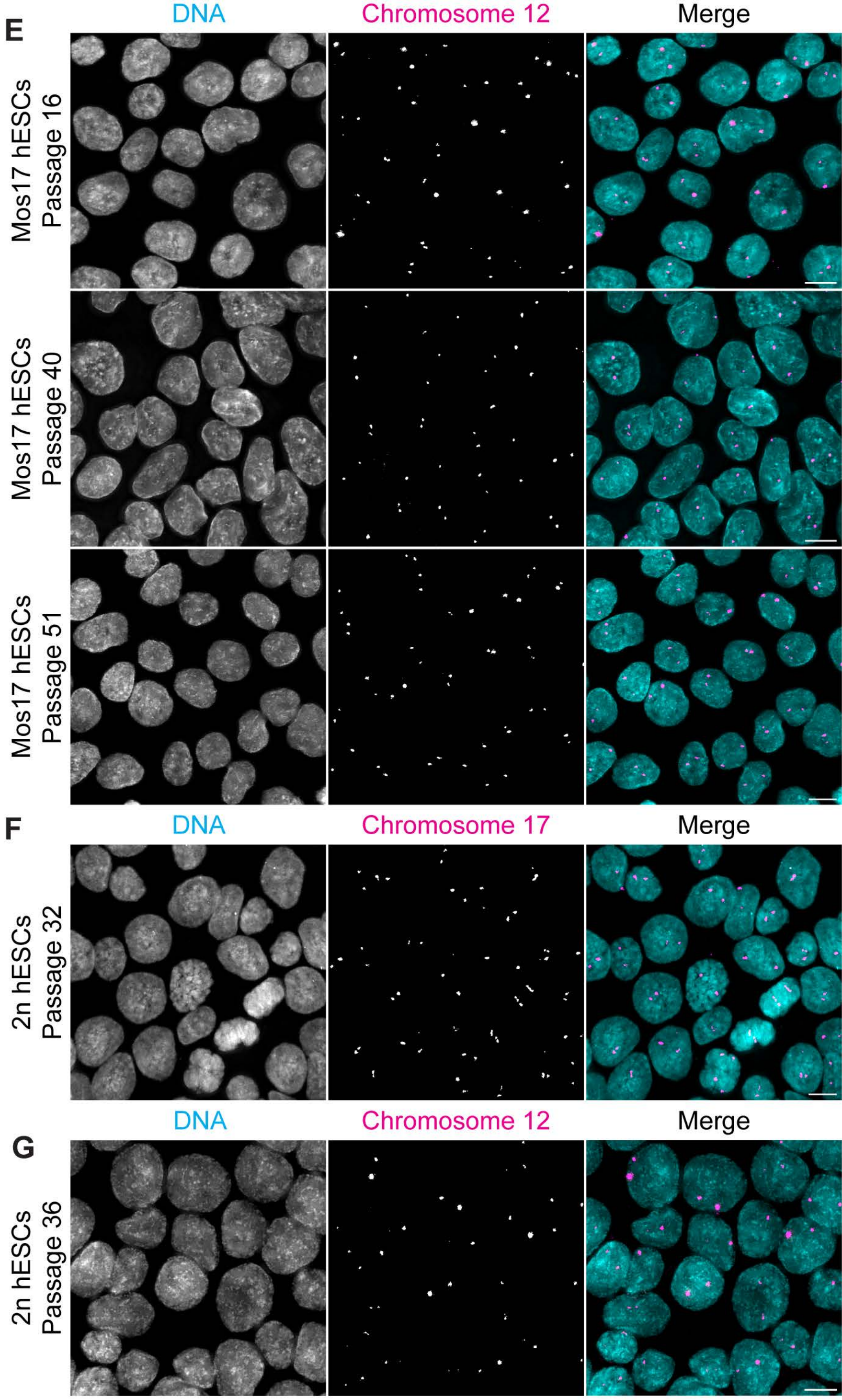


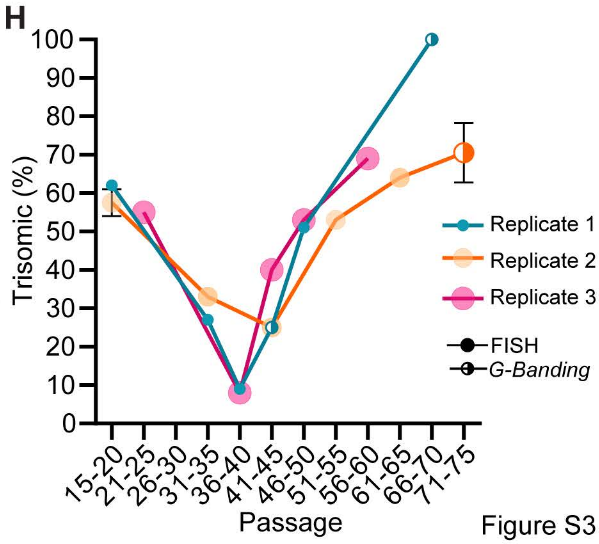


Figure S3

**Figure S3. STR analysis, karyotype analysis, FISH analysis for chromosomes 12 and 17 in 2n or Mos17 hESCs, and replicates for the percent of trisomic Mos17 hESCs. Related to Figure 3.** (**A**) Short tandem repeat (STR) analysis of Mos17 hESCs at passage 25 with approximately 50% disomic and trisomic chromosome 17 cells. (**B**) G-banding karyotype analysis of twenty 2n hESCs at passage 31 (left) and passage 57 (right). No clonal abnormalities were detected. (**C**) G-banding karyotype analysis of Ts12 hESCs at passage 36 (left) and passage 53 (right). At passage 36, all cells were trisomic for chromosome 12. At passage 53, twelve cells were trisomic for chromosome 12, three cells not shown were 47,XX,+12,der(21)t(1;21)(q12;p11.2), two cells not shown were 47,XX,dup(1)(q25q43),+12, and one cell disomic for chromosome 12. (**D**) G-banding karyotype analysis of twenty Mos17 hESCs at passage 20 (left), passage 30 (middle) and passage 69 (right). At passage 20, five cells were trisomic for chromosome 17, none at passage 30, and all at passage 69. (**E**-**G**) Representative images of chromosome 12 copy numbers in Mos17 hESCs (**E**) or of chromosome 17 (**F**) or 12 (**G**) copy numbers in 2n hESCs from FISH performed at different passages. Shown is DNA (cyan) and chromosome 17 or 12 (magenta). Scale bars: 10 μm. (**H**) Percent of Mos17 hESCs with trisomy 17 tracked by FISH (solid circle) or G-banding karyotype (half-filled circle) in three replicates labeled with different colors. Data points are binned into groups with 5 passages. $n \geq 100$ cells per bin and for bins with multiple data points per individual replicate the mean ± SD is shown.

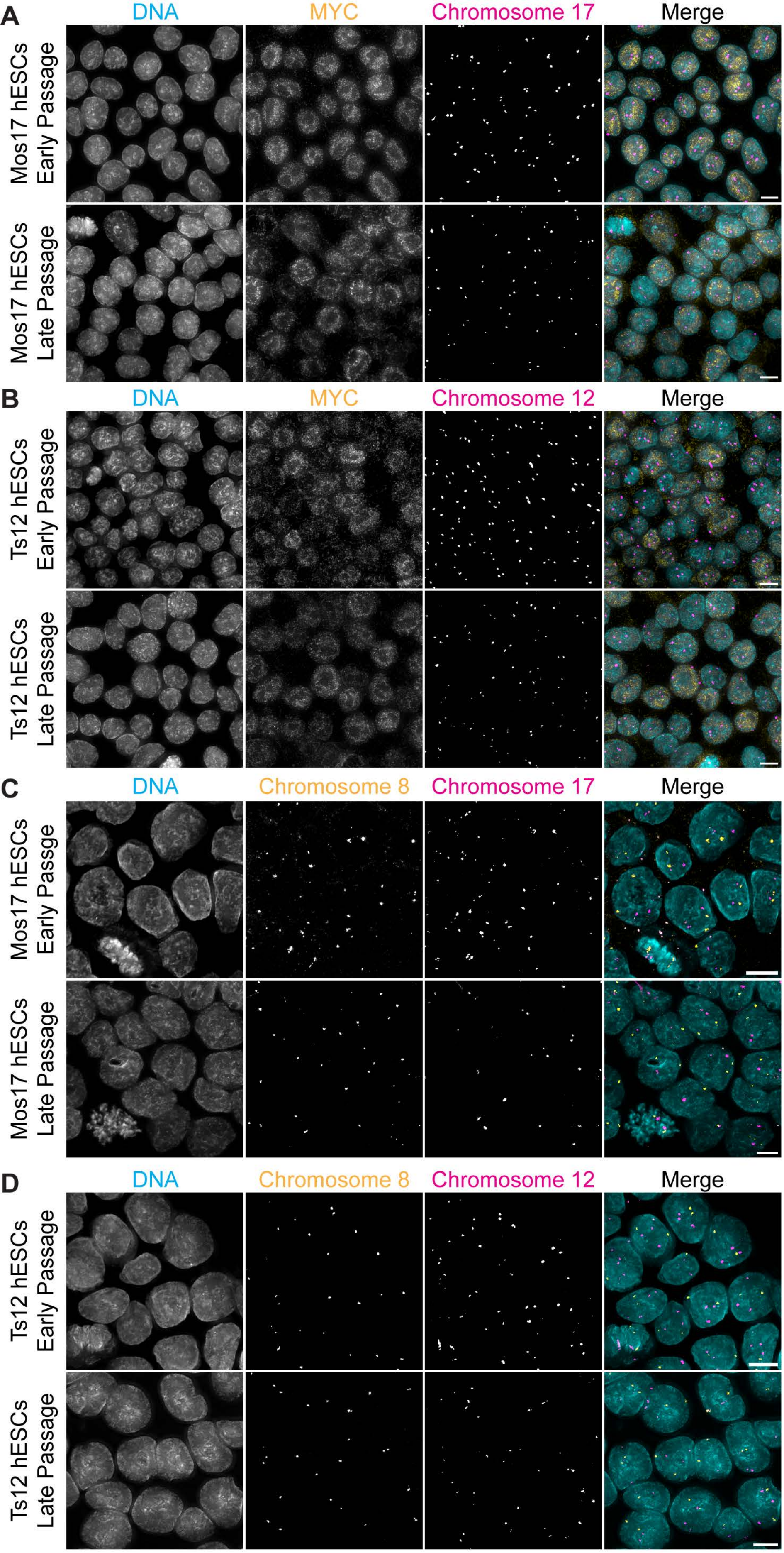


**E**

| **Mos17 hESCs** | | |
|---|---|---|
| Passage | Trisomic Chr 8 (%) | Disomic Chr 8 (%) |
| Early P:29 | ND | 100 |
| Late P:56 | ND | 100 |

**F**

| **Ts12 hESCs** | | |
|---|---|---|
| Passage | Trisomic Chr 8 (%) | Disomic Chr 8 (%) |
| Early P:45 | ND | 100 |
| Late P:65 | ND | 100 |

Figure S4

**Figure S4. Absolute Myc expression and chromosome 8 FISH analysis in Mos17 or Ts12 hESCs. Related to Figure 4.** (**A**-**B**) Representative IF-FISH images of Myc expression in a field of Mos17 (**A**) or Ts12 (**B**) hESCs at early (top) or late (bottom) passage. Shown is DNA (cyan), Myc (yellow) and chromosome 17 or 12 (magenta). Scale bars: 10 µm. (**C**-**D**) Representative images of chromosome 8 copy numbers in Mos17 hESCs (**C**) or Ts12 (**D**) hESCs from FISH performed at early (top) or late (bottom) passage. Shown is DNA (cyan), chromosome 8 (yellow), chromosome 17 or 12 (magenta) Scale bar: 5 µm. (**E**-**F**) Percent of Mos17 (**E**) or Ts12 (**F**) hESCs trisomic or disomic for chromosome 8 at early or late passage. One independent experiment; n = 100 cells per passage.

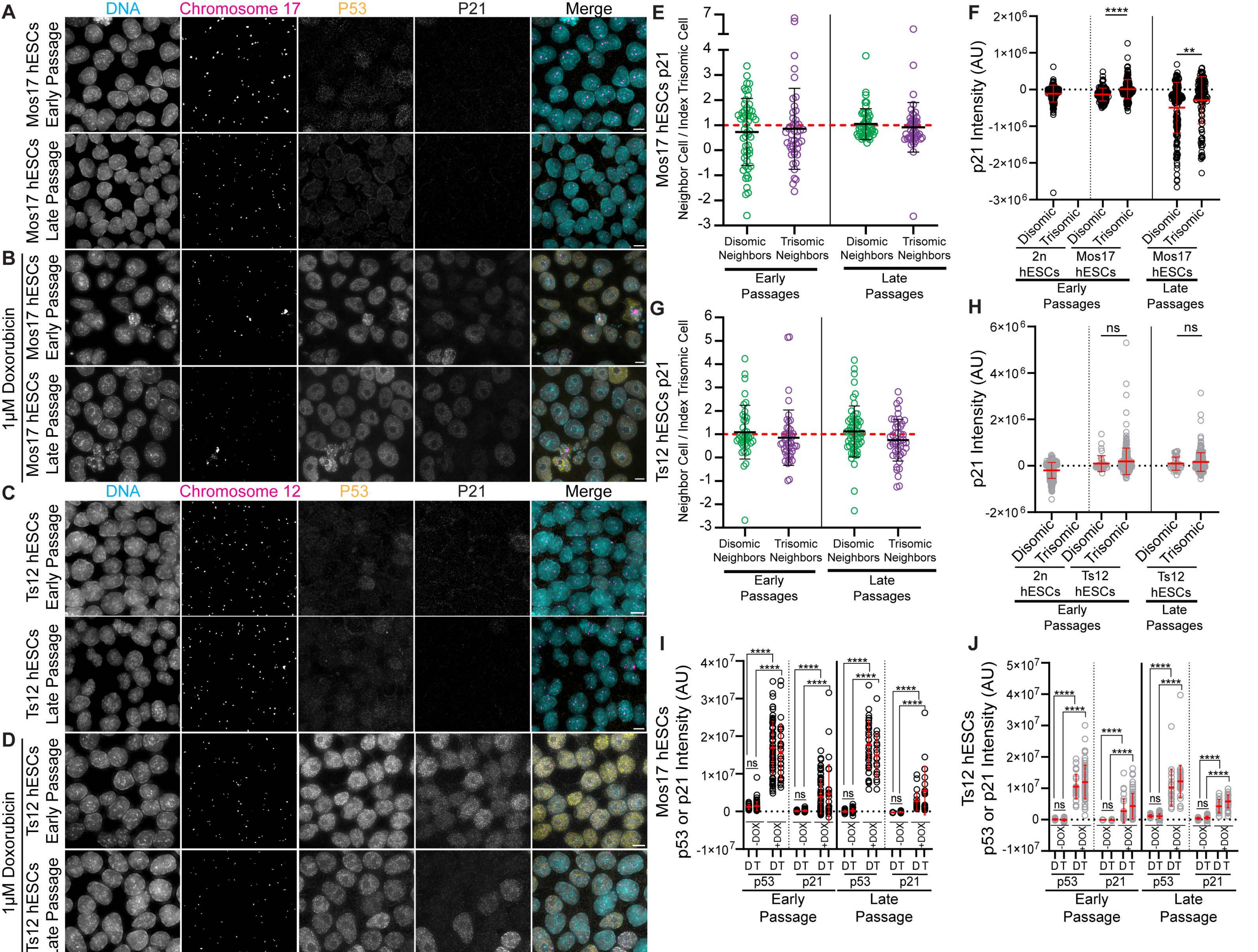
A
B
C
D
E
F
G
H
I
J
DNA
Chromosome 17
P53
P21
Merge
Mos17 hESCs Early Passage
Mos17 hESCs Late Passage
1µM Doxorubicin
Chromosome 12
Ts12 hESCs Early Passage
Ts12 hESCs Late Passage
Mos17 hESCs p21
Neighbor Cell / Index Trisomic Cell
Disomic Neighbors
Trisomic Neighbors
Early Passages
Late Passages
p21 Intensity (AU)
Disomic
Trisomic
2n hESCs
Mos17 hESCs
Ts12 hESCs
ns
Mos17 hESCs p53 or p21 Intensity (AU)
Ts12 hESCs p53 or p21 Intensity (AU)
-DOX
+DOX
D T
p53
p21
Early Passage
Late Passage
Figure S5

**Figure S5. p21 absolute and relative abundance levels and p53 and p21 quantification after DNA damage in Mos17, Ts12, or 2n hESCs. Related to Figure 5.** (**A**-**D**) Representative IF-FISH images for p53 and p21 basal expression or after induction of DNA damage with 1 μM doxorubicin for 4h in a field of Mos17 (**A**-**B**) or Ts12 (**C**-**D**) hESCs at early (top) or late (bottom) passage. Shown is DNA (cyan) chromosome 17 or 12 (magenta), p53 (yellow), and p21 (gray). Scale bars: 10 μm. (**E**) Quantification of p21 protein levels using IF-FISH. Neighbor cells were normalized to index trisomic cells in early (left) and late (right) passage Mos17 hESCs. Red dashed line indicates a ratio of 1 and equivalent p21 levels between index trisomic cells and neighbors. n = 60 index trisomic cells with > 100 neighbors; mean ± SD. (**F**) Quantification of p21 protein levels using IF-FISH in randomly selected, non-neighboring 2n and Mos17 hESCs at early (left panels) and late (right panels) passages. n= 300 total cells; mean ± SD. (**G**) Quantification of p21 protein levels using IF-FISH. Neighbor cells were normalized to index trisomic cells in early (left) and late (right) passage Ts12 hESCs. Red dashed line indicates a ratio of 1 and equivalent p21 levels between index trisomic cells and neighbors. n = 60 index trisomic cells with > 97 neighbors; mean ± SD. (**H**) Quantification of p21 protein levels using IF-FISH in randomly selected, non-neighboring 2n and Ts12 hESCs at early (left panels) and late (right panels) passages. n= 300 total cells; mean ± SD. (**I-J**) Quantification of p53 and p21 protein levels using IF-FISH in randomly selected, non-neighboring Mos17 (**I**) or Ts12 (**J**) hESCs after 4h treatment with 1 μM doxorubicin treatment at early (left panels) and late (right panels) passage. D = disomic, T = trisomic, n = 100 total cells; mean ± SD. Three (**E-H**) or one (**I-J**) independent experiment each for early and late passage; ns p > 0.05, **p < 0.01, or ****p < 0.0001 using a one sample t-test compared to hypothetical mean value 1 (**E** and **G**), a two-tailed Student's t-test for each cell line at early or late passage (**F** and **H**), or a one-way ANOVA and Tukey's multiple comparisons test (**I** and **J**).

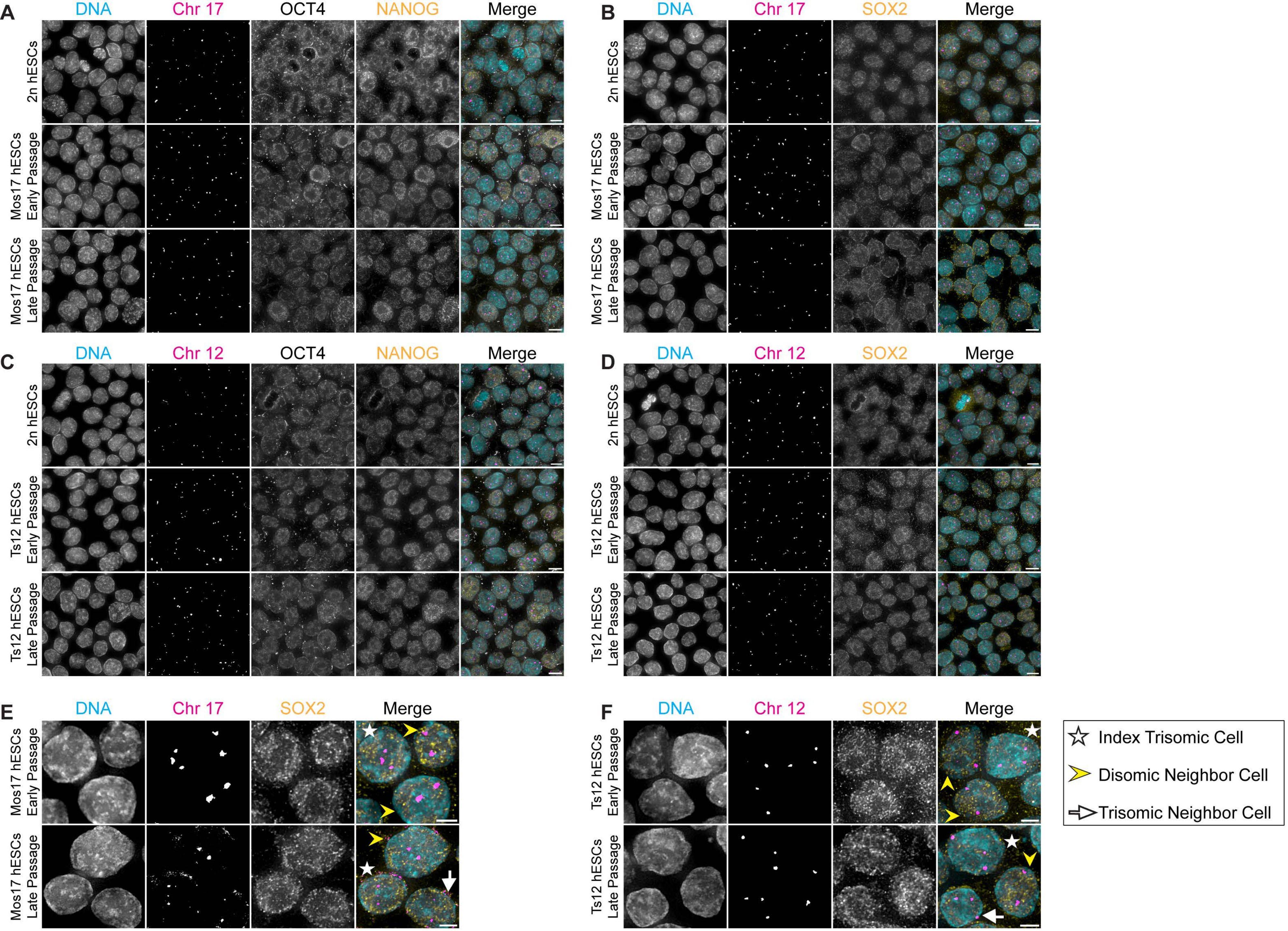
A
DNA
Chr 17
OCT4
NANOG
Merge
2n hESCs
Mos17 hESCs Early Passage
Mos17 hESCs Late Passage
B
DNA
Chr 17
SOX2
Merge
2n hESCs
Mos17 hESCs Early Passage
Mos17 hESCs Late Passage
C
DNA
Chr 12
OCT4
NANOG
Merge
2n hESCs
Ts12 hESCs Early Passage
Ts12 hESCs Late Passage
D
DNA
Chr 12
SOX2
Merge
2n hESCs
Ts12 hESCs Early Passage
Ts12 hESCs Late Passage
E
DNA
Chr 17
SOX2
Merge
Mos17 hESCs Early Passage
Mos17 hESCs Late Passage
F
DNA
Chr 12
SOX2
Merge
Ts12 hESCs Early Passage
Ts12 hESCs Late Passage
Index Trisomic Cell
Disomic Neighbor Cell
Trisomic Neighbor Cell

Figure S6

**Figure S6. Oct4, Nanog, and Sox2 absolute and relative expression in Mos17 or Ts12 hESCs.** (**A**-**D**) Representative IF-FISH images for Oct4, Nanog, or Sox2 expression in a field of 2n (**A**-**D**), Mos17 (**A**-**B**), or Ts12 (**C**-**D**) hESCs at early (middle panels) or late (bottom panels) passage. Shown is DNA (cyan), chromosome 17 or 12 (magenta), Oct4 (gray), and Nanog or Sox2 (yellow). Scale bars: 10 μm. (**E**-**F**) Representative images of Mos17 hESC (**E**) or Ts12 (**F**) neighborhoods comprised of an index trisomic cell (white star), disomic neighbor cells (yellow arrowhead), or trisomic neighbor cells (white arrow) at early (top) or late (bottom) passage. Show is DNA (cyan), chromosome 17 or 12 (magenta), and Sox2 (yellow). Scale bar: 5 μm.
